# AMPK-Fyn signaling promotes Notch1 stability to potentiate hypoxia-induced breast cancer stemness and drug resistance

**DOI:** 10.1101/458489

**Authors:** Mohini Lahiry, Saurav Kumar, Kishore Hari, Adithya Chedere, Mohit Kumar Jolly, Annapoorni Rangarajan

**Affiliations:** Department of Molecular Reproduction, Development and Genetics, Indian Institute of Science, Bangalore-560012, India; Centre for BioSystems Science and Engineering, Indian Institute of Science, Bangalore-560012, India

**Keywords:** Hypoxia, AMP-activated protein kinase (AMPK), Itch/AIP4, cleaved Notch1, breast cancer, cancer stem cell

## Abstract

Hypoxia is a hall mark of solid tumor microenvironment and contributes to tumor progression and therapy failure. The developmentally important Notch pathway is implicated in cellular response of cancer cells to hypoxia. Yet, the mechanisms that potentiate Notch signaling under hypoxia are not fully understood. Hypoxia is also a stimulus for AMP-activated protein kinase (AMPK), a major cellular energy sensor. In this study, we investigated if AMPK interacts with the Notch pathway and influences the hypoxia-response of breast cancer cells. Activating AMPK with pharmacological agent or genetic approaches led to an increase in the levels of cleaved Notch1 and elevated Notch signaling in invasive breast cancer cell lines. In contrast, inhibition or depletion of AMPK reduced the amount of cleaved Notch1. Significantly, we show that the hypoxia-induced increase in cleaved Notch1 levels requires AMPK activation. Probing into the mechanism, we demonstrate that AMPK activation impairs the interaction between cleaved Notch1 and its ubiquitin ligase, Itch/AIP4 through the tyrosine kinase Fyn. Under hypoxia, the AMPK-Fyn axis promotes inhibitory phosphorylation of Itch which abrogates its interaction with substrates, thus stabilizing cleaved Notch1 by reducing its ubiquitination and degradation. We further show that inhibition of AMPK alleviates the hypoxia-triggered, Notch-mediated stemness and drug resistance phenotype. Breast cancer patient samples also showed co-expression of hypoxia/AMPK/Notch gene signature. Our work thus establishes AMPK as a key component in the adaptation of breast cancer cells to hypoxia, and proposes therapeutic inhibition of AMPK to mitigate the hypoxia-triggered aggressiveness.

## Introduction

Developmental pathways such as Notch are known to regulate self-renewal and cell fate decisions in embryonic and tissue-specific stem cells (Artavanis-Tsakonas et al., 1999). In contrast, deregulation of Notch signaling is associated with malignant transformation (Radtke and Raj, 2003). Chromosomal translocation and activating mutations in Notch1 promote T-acute lymphoblastic leukemia (Aster et al., 2008). Aberrant Notch signaling has also been observed in several solid tumors including breast, cervical and prostate cancer (Allenspach et al., 2002). In breast cancer, we and others have reported an accumulation of intracellular domain of Notch1, as well as elevated expression of Notch ligand Jagged1 and its correlation with poor patient survival (Mittal et al., 2009; Reedijk et al., 2005; Stylianou et al., 2006). In contrast, there is loss of expression of negative regulators of Notch like Numb (Pece et al., 2004). Aberrant Notch signaling promotes cancer cell growth and survival, and alters metabolism. Recent studies have additionally implicated a critical involvement of Notch pathway in breast cancer stem cell self-renewal (D’Angelo et al., 2015) and therapy resistance (Wang et al., 2010).

Notch belongs to a family of four transmembrane receptors, named Notch 1-4 (Gordon et al., 2008). Ligand binding results in multiple proteolytic cleavages leading to the generation of cleaved Notch1 proteins, and finally the release of the transcriptionally active Notch intracellular domain (NICD) from the plasma membrane. NICD translocates to the nucleus where it forms a DNA-binding complex with other co-activators like MAML, CSL and p300, and activates expression of target genes like HES and HEY (Jarriault et al., 1995; Schroeter et al., 1998). Notch signaling is regulated at multiple levels, including presentation & availability of ligand, endocytosis & trafficking of processed receptors, and degradation by recognition of C-terminal PEST domain (Bray, 2006). In addition, post-translational modifications including phosphorylation, ubiquitination, hydroxylation and acetylation affect the stability, transcriptional activity and localization of NICD (Andersson et al., 2011).

Hypoxia develops in solid tumor environment due to rapid proliferation of cancer cells and an inadequate supply of oxygen to meet the demand. Under most scenarios, this hypoxia is likely to be cyclic in nature (Saxena and Jolly, 2019). Adaptation of tumor cells to hypoxia contributes to more aggressive and therapy-resistant phenotypes (Kim and Lee, 2017; Kunz and Ibrahim, 2003). Cellular adaptation to hypoxia is mainly orchestrated by the transcription factor HIF1-α (Wenger and Gassmann, 1997). The Hypoxia-HIF1 axis regulates cancer stemness by activating several pathways like TAZ (Xiang et al., 2014), CD47 (Zhang et al., 2015), PHGDH (Samanta et al., 2016), and ALKBH5 (Zhang et al., 2016). It also aids breast cancer cell migration and invasion through the Notch pathway (Sahlgren et al., 2008). Hypoxia potentiates Notch signaling by multiple mechanisms. It induces the expression of Notch ligands like Jagged2 and Delta-like ligand 4 (DLL4) (Jubb et al., 2009; Xing et al., 2011). Direct binding of HIF1-α to NICD leads to the upregulation of Notch-downstream genes (Gustafsson et al., 2005). In addition, the hypoxia-HIF1α axis also stabilizes NICD (Gustafsson et al., 2005); however, the downstream molecular mechanisms that lead to Notch stabilization remains poorly understood.

Yet another protein activated by hypoxia is the AMP-activated protein kinase, AMPK (Gusarova et al., 2011; Mungai et al., 2011). It is a heterotrimeric protein composed of catalytic α subunits, and regulatory β and γ subunits. AMPK is activated by stimuli that trigger an increase in AMP/ATP ratio, such as, glucose deprivation and chronic hypoxia (Hardie et al., 2012). Hypoxia is also known to activate AMPK independent of any changes in AMP/ATP ratio, through oxidant signaling (Emerling et al., 2009; Mungai et al., 2011).

Upon activation, AMPK inhibits anabolic pathways while activating catabolic pathways, thus bringing about energy homeostasis (Hardie et al., 2012). In addition to its central role in metabolism, recent studies have highlighted AMPK functions in regulating several physiological processes such as cell growth, polarity, apoptosis, autophagy and cell fate (Mihaylova and Shaw, 2011).

Owing largely to its growth suppressive effects, AMPK is mainly recognized for its tumor suppressive properties (Hardie and Alessi, 2013), and consistent with this notion, AMPK activating agents have been proposed for cancer treatment (Hardie, 2013). However, accumulating evidence points to a contextual oncogenic role of AMPK by enabling cancer cell survival under challenges faced in the tumor microenvironment, such as, glucose, oxygen, and matrix deprivation (Hindupur et al., 2014; Jeon et al., 2012; Jeon and Hay, 2012, 2015; Liang and Mills, 2013). In breast cancer, we and others have reported an increasing, grade-specific activation of AMPK in patient samples (Hart et al., 2015b; Sundararaman et al., 2016). We have also highlighted a novel role for AMPK in inhibiting apoptosis by phosphorylating PEA15 (Hindupur et al., 2014) and in mitigating the anchorage-deprivation stress by activating autophagy in breast cancer cells (Hindupur et al., 2014; Jeon et al., 2012; Saha et al., 2018). Hence, AMPK is increasingly recognized to mediate several non-metabolic functions by phosphorylating a plethora of proteins that participate in diverse cellular signaling pathways (Schaffer et al., 2015) which may be beneficial for tumor progression.

Thus, independent studies have demonstrated hypoxia-triggered activation of AMPK and Notch pathway; yet, a cross talk between these two pathways remains unexplored. In this study, we investigated if AMPK interacts with the Notch pathway and influences the response of breast cancer cells to hypoxia. We show that AMPK activation potentiates Notch signaling by stabilizing cleaved Notch1 through Fyn tyrosine kinase. We further show that inhibition or depletion of AMPK mitigates the hypoxia-driven, Notch1-mediated cancer stem cell and drug resistance phenotype. Our study thus highlights AMPK as a crucial component of the hypoxia-Notch1 signaling in promoting breast cancer aggressiveness.

## Results

### AMPK regulates cleaved Notch1 levels in breast cancer cells

To test if AMPK is involved in regulating Notch signaling, we first gauged the effect of pharmacologic and genetic activation of AMPK on Notch1 expression in breast cancer cell lines. To measure the Notch1 protein levels, we performed immunoblotting using an antibody specific to the C-terminus of Notch1 which recognizes both the full length (FL-Notch1; 300 kD) and proteolytically cleaved (cl-Notch1; ~120 kD) forms of Notch1. We used A769662 as a pharmacological activator of AMPK (Cool et al., 2006). Detection of an increase in phosphorylation of ACC, a bonafide substrate of AMPK (Winder and Hardie, 1996), confirmed AMPK activation upon treatment with A769662 **(Figure 1A)**. AMPK activation did not alter the transcript level of Notch1 **(Figure S1A)** or the full length Notch1 protein levels in MDA-MB-231 invasive breast cancer cell line **(Figure 1A)**; however, we observed elevated levels of cleaved Notch1 **(Figure 1A)**, suggesting possible activation of Notch signaling by AMPK.

**Figure 1:**
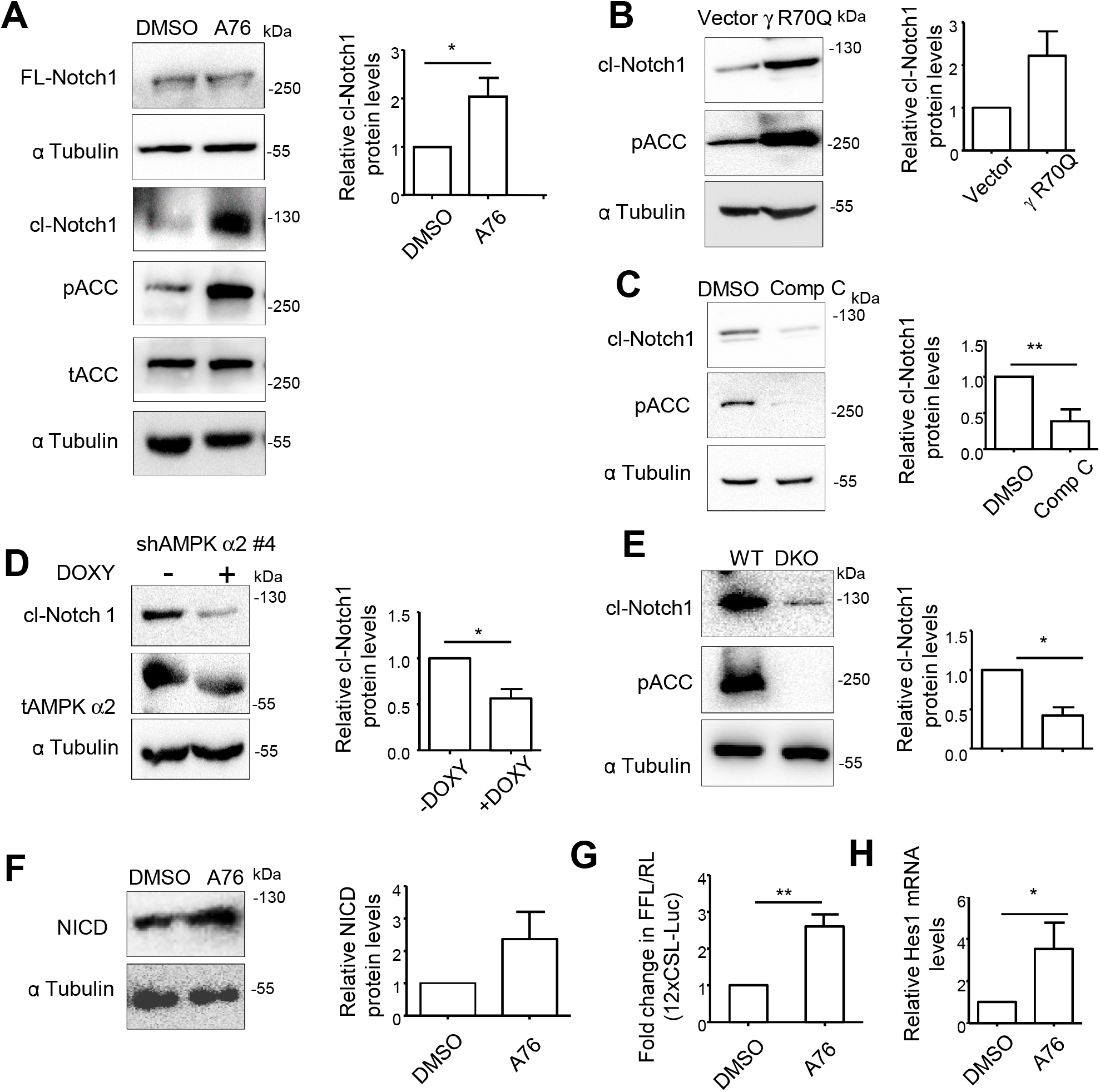
AMPK regulates cleaved Notch1 levels in breast cancer cells. (A) MDA-MB-231 cells were cultured in the presence of A769662 (A76; 100 μM) or DMSO as control and immunoblotted for indicated proteins. α Tubulin was used as loading control (n=3). Graphs in all immunoblotting experiments represent the densitometric analysis for quantification of relative amount of indicated protein and error bars represent ±SEM. (B) MDA-MB-231 cells transfected with constitutively active gamma subunit of AMPK (*γ* R70Q) or vector control and immunoblotted for indicated proteins (n=3). (C) MDA-MB-231 cells were treated with Compound C (Comp C; 10 μM) or DMSO as control for 24 h and immunoblotted for indicated proteins (n=6). (D) MDA-MB-231 cells stably expressing shAMPK α2 (#4) were induced with doxycycline (5μg/μl) for 48 h and immunoblotted for indicated proteins (n=4). (E) Immortalized wild type MEF and AMPKα1/2 DKO MEFs (DKO) were immunoblotted for indicated proteins (n=4). (F) MDA-MB-231 cells were cultured in the presence of A76 (100 μM) or DMSO as control for 24 h and subjected to immunoblotting for indicated protein (n=3). (G) MDA-MB-231 cells were transfected with 12xCSL-Luciferase (FFL) and Renilla TK (RL) and treated with A76 (100 μM) or DMSO as control for 48 h. Luciferase activity is represented as a ratio of Firefly (FFL) to Renila (RL) luciferase (n=4). (H) MDA-MB-231 cells were treated with A76 (100 μM) or DMSO as control for 48 h and RT-PCR analysis was carried out for *HES; GAPDH* served as housekeeping gene. Graph represents the fold change in *HES1* transcript levels normalized to *GAPDH* (n=4).

We next gauged the effect of AMPK activation on cleaved Notch1 levels in a variety of invasive, non-invasive and immortalized breast cells. We found elevated levels of cleaved Notch1 in invasive BT 474 and HCC 1806 breast cancer cells when treated with A769662 **(Figure S1B)**. However, in non-invasive MCF-7 and immortalized HMLE breast cells (Elenbaas et al., 2001), cleaved Notch1 levels were not modulated by AMPK activator **(Figure S1B)**, suggesting that the regulation of cleaved Notch1 protein levels by AMPK is likely to be cell-type and context-specific.

Since pharmacological agents can have non-specific effects, we additionally used genetic approaches to activate AMPK and checked for cleaved Notch1 levels. Over expression of constitutively activated AMPK with γR70Q mutation (Hamilton et al., 2001) led to increased AMPK activity, as gauged by increase in pACC levels, as well as higher levels of cleaved Notch1 **(Figure 1B)**. Similarly, overexpression of a constitutively active form of an AMPK upstream kinase CaMKKβ also resulted in elevated cleaved Notch1 levels **(Figure S1C)**. Thus, increasing AMPK activity led to elevated cleaved Notch1 levels in invasive breast cancer cells.

To further address the role of AMPK in regulating cleaved Notch1 levels, we investigated the effects of AMPK inhibition and depletion using pharmacologic and genetic approaches. Upon treatment with a pharmacological inhibitor of AMPK, Compound C (Zhou et al., 2001), we observed reduced phosphorylated ACC levels (confirming the efficacy of the inhibitor) along with reduced levels of cleaved Notch1 **(Figure 1C)**. To further confirm, this effect using a genetic approach, we employed doxycycline-inducible knockdown strategy targeting the catalytic α subunit of AMPK. In keeping with the higher expression and activity of AMPKα2 isoform, compared to AMPKα1, in breast cancer cell lines (Hindupur et al., 2014), we gauged the effect of AMPKα2 knockdown on cleaved Notch1 levels. Addition of doxycycline to MDA-MB-231 cells stably expressing the inducible shAMPKα2 (clone#4) construct led to a reduction in the levels of total AMPK α2, as well as a reduction in the levels of cleaved Notch1, compared to un-induced control cells **(Figure 1D)**. AMPK depletion using an independent, doxycycline-inducible, shRNA construct (clone#1) against AMPKα2 also led to a reduction in cleaved Notch1 levels **(Figure S1D)**. To further confirm the role of AMPK, we used AMPK α1/α2 double knock out (AMPK-DKO) MEFs. Compared to wild type (WT) MEFs, AMPK-DKO MEFs showed reduced levels of cleaved Notch1 **(Figure 1E)**. Restoration of AMPK expression in DKO MEFs by exogenous expression of catalytic subunits of AMPK led to increase in cleaved Notch1 levels **(Figure S1E)**, further confirming a direct role for AMPK in the positive regulation of cleaved Notch1 levels.

### AMPK activation promotes Notch signaling

We next investigated whether an increase in cleaved Notch1 levels upon AMPK activation led to higher Notch downstream signaling. To do so, we first checked if AMPK activation leads to elevated levels of the active form of cleaved Notch1. The C-terminal specific antibody that we have used is likely to recognize both cleaved but membrane tethered form of Notch1, as well as the active intracellular domain (ICD) of Notch1 that translocates to the nucleus (Brou et al., 2000). Cleavage by gamma-secretase at Valine 1744 results in the generation of active form of Notch1 (NICD) which enhances transcription of downstream genes. We therefore checked the effect of AMPK activation in the generation of NICD using an antibody that specifically recognizes the gamma secretase-specific cleaved Notch1 (Valine 1744). Immunoblotting revealed elevated levels of NICD in the presence of AMPK activator A769662 **(Figure 1F)**. To further confirm a role for AMPK in modulating NICD levels, we tested the effect of AMPK knockdown in cells ectopically expressing NICD. We chose HEK293T cells for this experiment as they show very minimal basal levels of endogenous Notch1 expression. AMPK depletion led to reduction in the levels of exogenously expressed NICD **(Figure S1F)**, additionally suggesting that the effect of AMPK on accumulation of cleaved Notch1 may be downstream of gamma secretase cleavage, perhaps by altering NICD stability.

In light of our data revealing an increase in active, cleaved Notch1 levels upon AMPK activation, we next tested if AMPK modulates Notch signaling. To do so, we used the Notch-responsive 12xCSL-luciferase reporter assay (Blokzijl et al., 2003). Treatment with AMPK activator A769662 led to a 2.6-fold increase in 12xCSL-luciferase reporter activity **(Figure 1G)**, indicative of increase in canonical Notch signaling upon AMPK activation. To further gauge the effect of AMPK activation on Notch signaling, we measured the endogenous levels of Notch target genes. Treatment with AMPK activator also led to an increase in the levels of two Notch pathway downstream genes Hes1 **(Figure 1H)** and Hes5 **(Figure S1G)**, as gauged by RT-PCR and immunocytochemistry, respectively. In addition, a microarray-based data of A769662-treated MDA-MB-231 cells revealed upregulation of several Notch pathway-regulated genes including HES5, HEY2, SNAI2, HEYL, IFNG and CDH6 **(Figure S1H)**. Taken together, these observations reveal that elevated cleaved Notch1 levels downstream to AMPK activation leads to enhanced Notch signaling.

### AMPK inhibition or depletion impairs hypoxia-induced NICD generation

In response to hypoxia, independent reports have shown elevated Notch signaling in breast cancer (Sahlgren et al., 2008) and AMPK activation in other cell types (Emerling et al., 2009; Mungai et al., 2011). In light of our data revealing elevated Notch signaling upon AMPK activation in breast cancer cells, we investigated if AMPK is involved in hypoxia-induced Notch signaling. To address this, we treated MDA-MB-231 cells with a hypoxia-mimetic agent CoCl_2_, which simulates hypoxic effects by stabilizing HIF-1α (Xi et al., 2004). As shown before (Chen et al., 2017), treatment with CoCl_2_ led to an increase in phosphorylated AMPK levels in MDA MB 231 cells **(Figure 2A)**. It also led to an increase in cleaved Notch1 levels **(Figure 2A)**. Treatment of yet another invasive breast cancer cell line BT474 with CoCl_2_ led to increase in carbonic anhydrase IX (CA9), which is a direct readout of HIF-1α, and pACC levels, which is an indicator of AMPK activity **(Figure S2A)**. Consistent with our previous results, we observed elevated levels of cleaved Notch1 **(Figure S2A)**. Having confirmed both AMPK activation and elevated cleaved Notch1 levels in the presence of CoCl_2_, we investigated if AMPK is involved in hypoxia-triggered elevated cleaved Notch1 levels. To do so, we tested the effect of AMPK inhibition in this process under hypoxia. CoCl_2_-induced elevated cleaved Notch1 was impaired in the presence of AMPK inhibitor, Compound C **(Figure 2B)**. Similar results were observed in BT-474 cells also **(Figure S2A)**. To further confirm this, we measured the effect of AMPK depletion in the CoCl_2_-induced cleaved Notch 1 levels. Cells expressing the control pTRIPZ construct showed expected increase in cleaved Notch1 levels in the presence of CoCl_2_ when treated with doxycycline, whereas those expressing inducible AMPKα2 shRNA construct (clone# 1) failed to do so **(Figure 2C)**. We obtained similar results with an independent inducible AMPKα2 (clone#4) shRNA construct **(Figure S2B)**. Together, these data indicated a role for AMPK in the CoCl_2_-induced increase in cleaved Notch1 levels.

**Figure 2:**
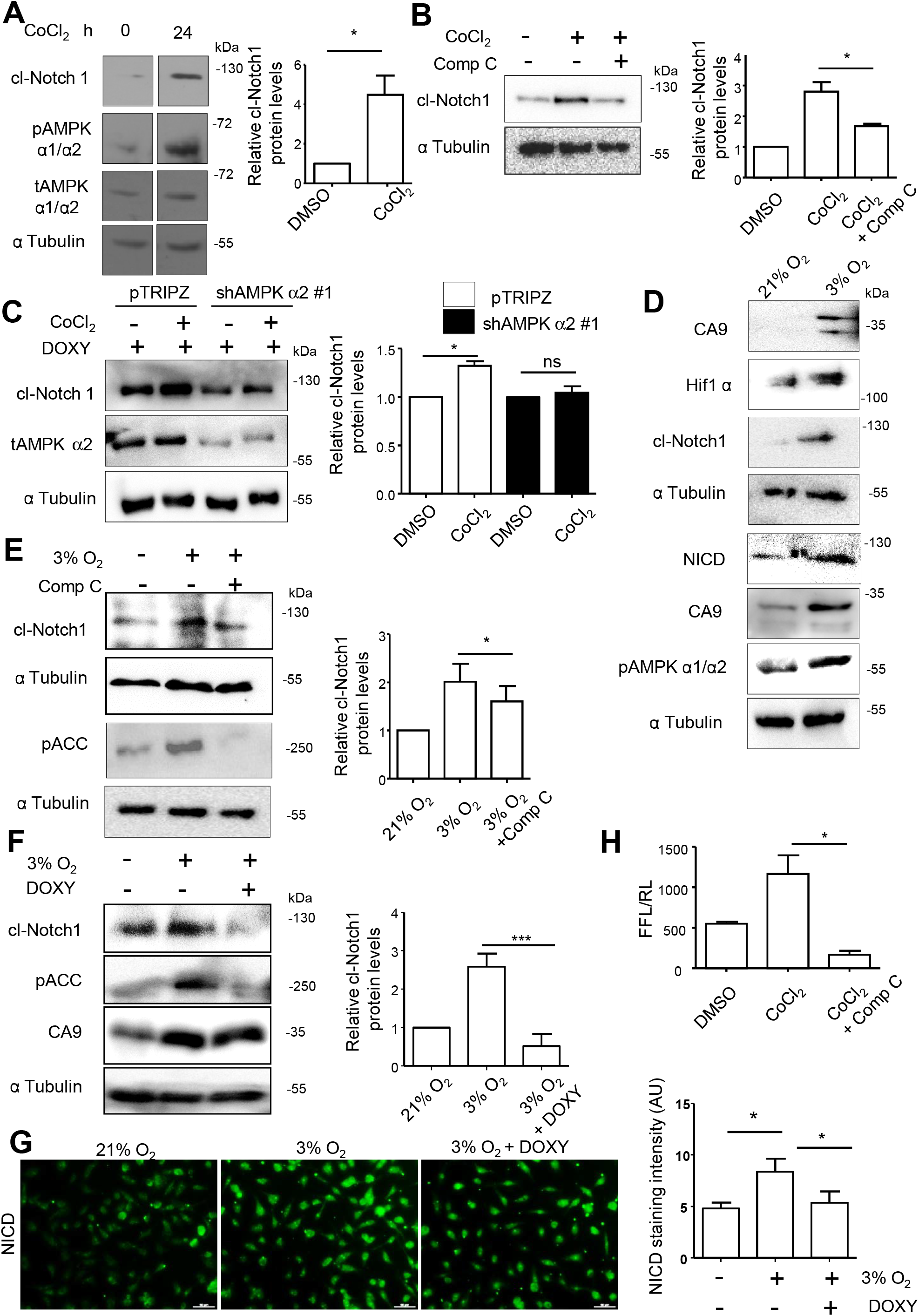
AMPK inhibition or knockdown impairs hypoxia-induced cleaved Notch1 generation. (A) MDA-MB-231 cells were subjected to CoCl_2_ (150 μM) treatment for 24 h and immunoblotted for indicated protein; lanes from the same run were assembled together. Graphs in all immunoblots represent the densitometric analysis for quantification of relative amount of indicated protein (n=4). (B) MDA-MB-231 cells were subjected to CoCl_2_ (150 μM) with DMSO or Comp C (10 μM) treatment as indicated for 24 h and immunoblotted for indicated proteins (n=4). (C) MDA-MB-231 cells stably expressing shAMPK α2 (#1) and vector control pTRIPZ were induced with doxycycline (5μg/μl) for 24 h followed by treatment with CoCl_2_ (150 μM) and immunoblotted for indicated proteins (n=3). (D) MDA-MB-231 cells were grown in 21% or 3% O_2_ for 24 h and harvested for immunoblotting for indicated proteins. (E) MDA-MB-231 cells were grown in 21% or in 3% O_2_ in the presence or absence of Comp C (10 μM) and immunoblotted for indicated proteins (n=3). (F) MDA-MB-231 cells stably expressing shAMPK α2 (#4) were induced with doxycycline (5μg/μl) for 24 h and subsequently grown in 3% O_2_ and immunoblotted for indicated proteins (n=3). (G) MDA-MB-231 cells stably expressing shRNA against AMPK α2 (#4) were induced with doxycycline (5μg/μl) for 24 h and subsequently grown in 3% O_2_ for 24 h and were immunostained for valine 1744 cleavage-specific NICD protein (Scale bar: 50 μm) (n=3). Graph represents the mean intensity of valine 1744 specific NICD protein expression in 3 independent experiments. (H) HEK-293T cells were transfected with 12xCSL-Luciferase (FFL) and Renilla TK (RL) and NICD expression construct. The cells were treated with CoCl_2_ for 48 h with DMSO or Compound C. Luciferase activity is represented as a ratio of Firefly (FFL) to Renila (RL) luciferase (n=3). Error bars represent ±SEM in all graphs.

To better study the AMPK-Notch crosstalk under hypoxia relevant to physiological and patho-physiological condition, we used oxygen-regulatable trigas incubator to create hypoxic 3% oxygen condition in contrast to ambient 21% oxygen in regular incubators. Cells cultured in hypoxia showed increased levels of HIF-1α and CA 9, indicative of successful generation of hypoxic condition **(Figure 2D)**. As expected, we also observed elevated levels of cleaved Notch1 and pAMPK under hypoxia compared to normoxia **(Figure 2D)**. However, AMPK inhibition with Compound C or depletion of AMPKα2 impaired the hypoxia-induced increase in cleaved Notch1 levels **(Figure 2E-F)**, thus confirming a role for AMPK in the hypoxia-induced elevated cleaved Notch1 levels.

To further probe if AMPK plays a role in the generation of the active form of cleaved Notch1 under hypoxia, we used the gamma secretase cleavage specific Notch1 (Valine 1744) antibody. We observed elevated NICD levels under hypoxia by immunoblotting **(Figure 2D)**. In addition, we performed immunocytochemistry to gauge NICD levels. As expected, we observed an increase in pAMPK staining intensity under hypoxia, confirming the generation of hypoxic condition. We also observed an increase in NICD staining intensity under hypoxia compared to cells grown in normoxia **(Figure S2C)**. Further, AMPK inhibition reduced the hypoxia-triggered NICD staining intensity **(Figure 2G)**, suggesting a role for AMPK in the generation of active, cleaved form of Notch1 in hypoxia.

We next asked if the positive regulation of cleaved Notch1 levels by AMPK in response to hypoxic condition is also reflected in Notch signaling. To test this, we probed the activity of the Notch-responsive 12XCSL-reporter construct in the presence of CoCl_2_ to mimic hypoxic condition, and tested the role of AMPK. As expected, treatment with CoCl_2_ increased the activity of the 12XCSL-reporter; however, inhibition of AMPK impaired this **(Figure 2H)**. Taken together, these results reveal a key role for AMPK in the accumulation of active, cleaved Notch 1 and positive regulation of the Notch signaling pathway in response to hypoxic stress.

### AMPK inhibition affects stability of cleaved Notch1 under hypoxia

In light of our observation that AMPK activation increases the amounts of cleaved Notch1, but not full-length receptor, we next explored if AMPK enhances the stability of cleaved Notch1. To do so, we performed a cycloheximide chase assay in MDA-MB-231 cells harboring doxycycline-inducible AMPKα2 shRNA. Addition of cycloheximide revealed a faster reduction in the amount of cleaved Notch1 under AMPK depleted conditions **(Figure 3A)**, suggesting a possible role for AMPK in regulating cleaved Notch1 stability. To further confirm this, we subjected AMPK DKO MEFs to cycloheximide treatment. Treatment with cycloheximide revealed a higher reduction in cleaved Notch1 levels in AMPK DKO MEFs compared to WT MEFs **(Figure S3A)**. Together, these data showed that the absence of AMPK enhances the rate of Notch1 degradation, suggesting a role for AMPK in stabilization of cleaved Notch1.

**Figure 3:**
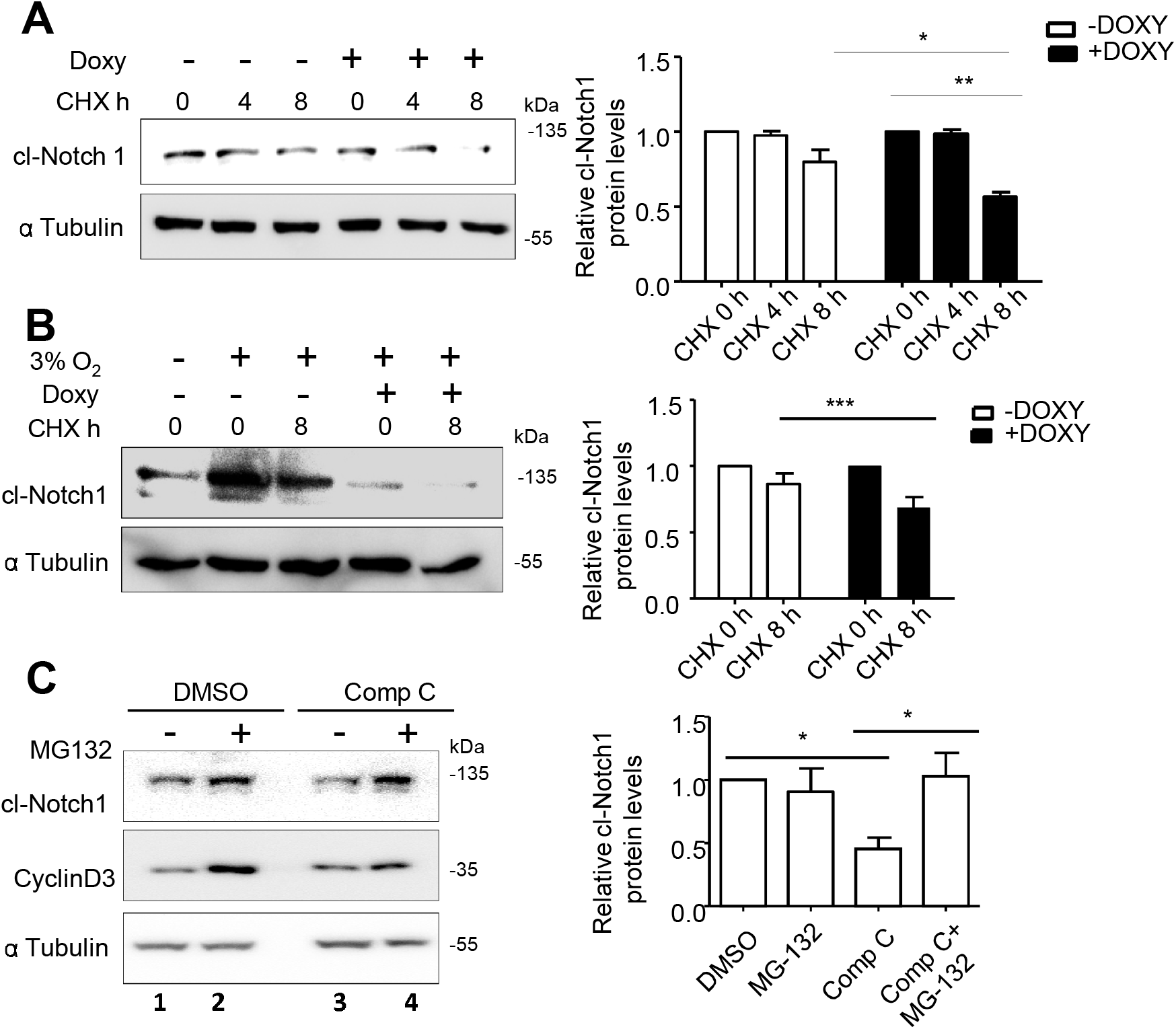
AMPK inhibition affects stability of cleaved Notch1 under hypoxia. (A-B) MDA-MB-231 cells stably expressing shAMPK α2 (#4) were induced by doxycycline (5μg/μl) for 48 h and (A) subsequently treated with cycloheximide for 4 h and 8 h and immunoblotted for indicated proteins; and graphs represent the densitometric analysis for quantification of relative amount of indicated protein and (B) grown in 3% O_2_ for another 24 h followed by cycloheximide treatment for 8 h and immunoblotted for indicated; (n=3 each). (C) MDA-MB-231 cells were treated with Comp C (10 μM) or DMSO control for 24 h and subsequently treated with or without MG-132 (5 μM) for 8 h. Cells were immunoblotted for indicated proteins (n=3); Error bars represent ±SEM in all graphs.

We next investigated if AMPK is involved in the hypoxia-induced stabilization of cleaved Notch1 as shown earlier **(Figure 2)**, and as reported previously in P19 cells (Gustafsson et al., 2005). To do so, we grew MDA-MB-231 cells harbouring doxycycline-inducible AMPKα2 shRNA constructs in trigas incubator with 3% oxygen and subsequently treated with cycloheximide. Immunoblotting revealed heightened reduction of cleaved Notch1 levels under hypoxia in the presence of doxycycline **(Figure 3B)**. Similarly, addition of AMPK inhibitor Compound C to MDA-MB-231 cells exposed to CoCl_2_ also revealed heightened reduction in cleaved Notch1 levels compared to vehicle control DMSO **(Figure S3B)**. Together, these data confirmed a key role for AMPK in regulating cleaved Notch1 stability in response to hypoxia.

Since cleaved Notch1 is reported to be targeted for degradation through the 26S proteasomal machinery (Gupta-Rossi et al., 2002), we next investigated if AMPK affects this process. To address this, we used the proteasomal complex inhibitor MG132, and asked if reduced cleaved Notch1 levels observed under AMPK inhibited condition can be rescued by proteasomal inhibition. As expected, treatment with MG132 led to stabilization of cyclinD3, which served as a positive control (lane 2 vs 1; **Figure 3C**). Further, we found that Compound C-mediated reduction in cleaved Notch1 levels (lane 3 vs 1) was rescued in the presence of MG132 (lane 4 vs 3; **Figure 3C**). Thus, AMPK activation plays a critical role in cleaved Notch1 stabilization by preventing its degradation through the proteasomal complex.

### AMPK inhibits Notch1 ubiquitination by modulating its interaction with Itch

Since protein degradation by the proteasomal system involves tagging of target proteins with ubiquitin, we next investigated if AMPK stabilizes cleaved Notch1 by regulating its ubiquitination. To address this question, we first asked if AMPK alters Notch1 K48-linked polyubiquitination which specifically marks proteins for proteasomal degradation. To do so, we immunoprecipitated Notch1 and probed by immunoblotting using an antibody specific for K48-linked polyubiquitin. We observed a reduction in K48-linked ubiquitination of cleaved Notch1 in the presence of AMPK activator A769662 **(Figure 4A)**. In the reverse experiment, when we inhibited AMPK using Compound C, the K48-linked ubiquitination of cleaved Notch1 was elevated **(Figure S4A)**. These data suggested that AMPK possibly stabilizes cleaved Notch1 by regulating its ubiquitination, and thereby degradation by the proteasomal pathway.

**Figure 4:**
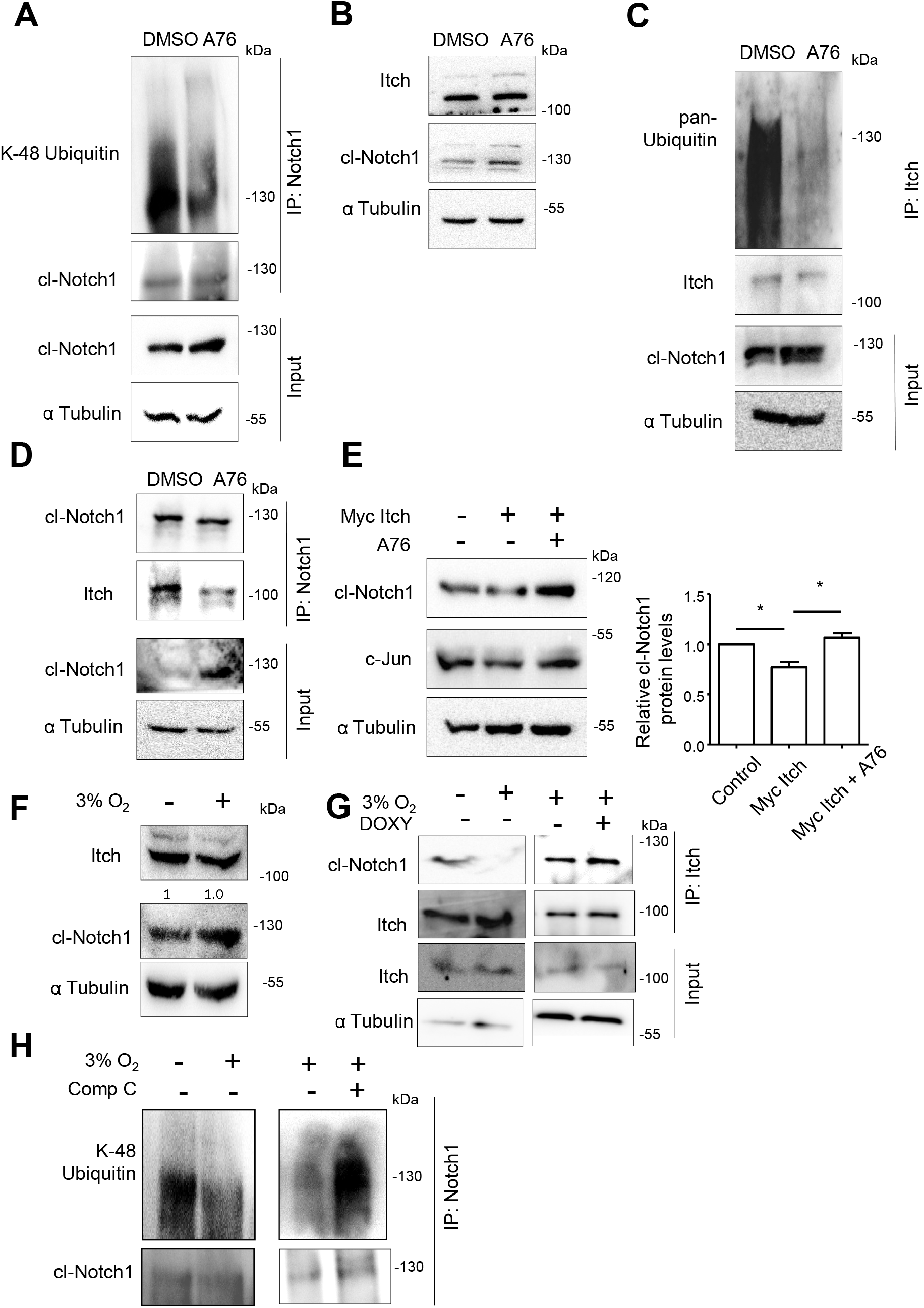
AMPK inhibits Notch1 ubiquitination by modulating its interaction with Itch. (A) MDA-MB-231 cells were cultured in the presence of A76 (100 μM) or DMSO control for 24 h and subsequently treated with MG132 (5 μM) for 2 h prior to Notch1 immuno-precipitation using the C-terminus specific Notch1 antibody followed by immunoblotting for anti-K-48 ubiquitin linkage specific antibody (n=3). (B) MDA-MB-231 cells were cultured in the presence of A76 (100 μM) or DMSO control for 24 h and subsequently immunoblotted for Itch protein (n=3). (C) MDA-MB-231 cells were cultured in the presence of A76 (100 μM) or DMSO control for 24 h and subsequently treated with MG132 (5 μM) for 2 h prior to Itch immuno-precipitation and immunoblotted for pan-ubiquitin. (D) MDA-MB-231 cells were cultured in the presence of A76 (100 μM) or DMSO control for 24 h and subsequently treated with MG132 (5 μM) for 2 h prior to Notch1 immuno-precipitation and immunoblotted for Itch protein (n=3). (E) MDA-MB-231 cells were transfected with vector control or Myc tagged Itch plasmid. Cells were cultured in the presence of A76 (100 μM) or DMSO control and immunoblotted for indicated proteins. Graph represents the densitometric analysis for quantification of relative amount of indicated protein (n=3). (F) MDA-MB-231 cells grown in 21% O_2_ or 3% O_2_ were immunoblotted for indicated proteins (n=3). (G) MDA-MB-231 cells stably expressing shAMPK α2 (#1) were grown for 24 h in 21% O_2_ or 3% O_2_ with and without induction by doxycycline (5μg/μl) (as indicated) and Itch was immuno-precipitated and immunoblotted for Notch1 protein. Same lysates were used for both the panels. Immuno-precipitation was carried out separately for the 2 panels and immunoblotted separately. (n=3) (H) MDA-MB-231 cells grown in 21% O_2_ or 3% O_2_ were treated with DMSO or Comp C (10 μM) for 24 h and with MG132 (5 μM) for 2 h prior to harvesting. Cleaved Notch1 was immuno-precipitated followed by immunoblotting using K-48 linked ubiquitin-specific antibodies (n=3). Lysates were prepared separately and immuno-precipitation and immunoblotting were carried out separately for the 2 panels.

To further explore the mechanism downstream of AMPK in mediating cleaved Notch1 stability, we investigated into post-translational modifications, such as, Ser/Thr phosphorylation and acetylation that are known to stabilize Notch1 protein levels (Andersson et al., 2011). We did not detect much change in the overall levels of Ser/Thr phosphorylation on immuno-precipitated cleaved Notch1 upon AMPK activation **(Figure S4B)**. Further, bioinformatic analysis revealed Notch1 T1998 as a putative AMPK phosphorylation site. To test if this site is involved in cleaved Notch1 stability, we carried out site-directed mutagenesis using an NICD expressing construct at threonine 1998 and changed it to non-phosphorylatable alanine (T1998A). If cleaved Notch1 stability is mediated by phosphorylation of this site, we hypothesized that the non-phosphorylatable alanine mutant would be insensitive to AMPK activation, and thus undergo increased degradation. To investigate this, wild type NICD and an engineered NICD harbouring T1998A substitution were transfected into HEK 293T cells, treated with A769662, and cleaved Notch1 levels were gauged. NICD T1998A responded to AMPK activation in a manner similar to wildtype NICD **(Figure S4C)**. This indicated that the effect mediated by AMPK is not through phosphorylation of Notch1 at residue T1998. Further, using an acetyl lysine specific antibody, we also failed to detect much change in the overall acetylation status of cleaved Notch1 upon AMPK activation **(Figure S4D)**.

Since we failed to observe changes in the overall Ser/Thr phosphorylation or acetylation status of cleaved Notch1 in response to AMPK activation, we shifted our focus to ubiquitin ligases. Notch1 is known to be a substrate for several ubiquitin ligases such as FBXW-7, Itch/AIP4, NEDD 4, Deltex 1, c-Cbl (Moretti and Brou, 2013). Notably, a study showed that AMPK activation disrupts the interaction of Itch/AIP4 (henceforth referred to as Itch) with its substrate p73 (Adamovich et al., 2014). We therefore examined a possible role for Itch in AMPK-mediated cleaved Notch1 stability. To do so we first examined the levels of Itch upon AMPK activation. We failed to see changes in the levels of Itch in the presence of AMPK activator A769662 **(Figure 4B)**. We then queried the activity of Itch by checking its auto-ubiquitination status. Itch auto-ubiquitinates itself by the non-degradative, K63 linkages (Scialpi et al., 2008), which also positively regulates its activity. We immunoprecipitated Itch and checked for its ubiquitination status by immunoblotting. Immunoprecipitated endogenous Itch revealed highly diminished ubiquitin smear in the presence of AMPK activator A769662 compared to DMSO vehicle control **(Figure 4C)**. We next investigated if AMPK activation affects the Itch-Notch interaction. Indeed, immunoprecipitation of Notch1 followed by immunoblotting with Itch revealed a reduced interaction between Notch1 and Itch in the presence of AMPK activator A769662 **(Figure 4D)**. These data suggested a possible mechanism downstream of AMPK activation leading to cleaved Notch1 stability by regulating Itch-Notch1 interaction.

To further confirm a role for AMPK in Itch-mediated regulation of cleaved Notch 1 stability, we asked if AMPK activation would rescue the effects of Itch overexpression on cleaved Notch1 levels. As expected, Itch overexpression led to a decrease in the amounts of another known substrate of Itch, c-Jun, as well as cleaved Notch1, whereas the presence of AMPK activator A769662 impaired this effect **(Figure 4E)**. Overexpression of Itch in yet another invasive breast cancer cell line, BT-474, yielded similar down-modulation of cleaved Notch1 which was rescued on AMPK activation **(Figure S4E)**. Taken together, these observations suggested a protective role for AMPK in Itch-mediated Notch degradation.

We next asked if AMPK plays a similar role in cleaved Notch1 stabilization in response to hypoxia by modulating Itch-Notch interaction. To address this, we first investigated Itch levels under hypoxia, and found that Itch levels remained unchanged in hypoxic condition of 3% O_2_ compared to normoxia **(Figure 4F)**. We next gauged Notch-Itch interaction in hypoxia, as well as tested the effect of AMPK depletion in this interaction. To do so, we used MDA-MB-231 cells harboring the doxycycline inducible AMPK shRNA system. To gauge Itch-Notch interaction, we immunoprecipitated endogenous Itch and measured the amounts of associated Notch1 by immunoblotting. Under un-induced (minus doxycycline) condition, we observed a reduction in cleaved Notch1 levels associated with immunoprecipitated Itch in hypoxia compared to normoxia (lanes 2 vs 1; **Figure 4G**). Upon induction of AMPK depletion with doxycycline, we observed higher amount of cleaved Notch1 in the Itch-immunoprecipitate under hypoxia (lane 4 vs 3), suggesting higher Itch-Notch interaction upon AMPK depletion. These data supported a role for AMPK in regulating the Itch-Notch1 interaction under hypoxia.

We previously showed a direct role for AMPK in regulating the K-48 linked degradative ubiquitination of Notch1 (**Figure 4A** and **Figure S4A**). We next inquired if the regulation of cleaved Notch1-Itch interaction under hypoxia affects the K-48 linked degradative ubiquitination of cleaved Notch1 in an AMPK-dependent manner. To address this, we performed immunoprecipitation of Notch1 under hypoxia in the presence and absence of AMPK inhibitor Compound C, and probed with K-48 linkage specific anti-ubiquitin antibodies. We found a reduction in the Notch1 K-48 associated ubiquitination in hypoxia **(Figure 4H)**. In contrast, AMPK inhibition increased the K-48 linked ubiquitination of cleaved Notch1 in hypoxic conditions **(Figure 4H)**. Thus, these data uncover a novel AMPK-mediated regulation of cleaved Notch1 stability in hypoxia through modulating its interaction with the ubiquitin ligase Itch.

### AMPK mediates Itch phosphorylation in hypoxia

We next sought to understand how AMPK negatively regulates the Itch-Notch interaction. Phosphorylation of Itch has been shown to regulate its interaction with substrates (Yang et al., 2006). In contrast to Ser/Thr phosphorylation, tyrosine phosphorylation of Itch negatively modulates its activity towards substrates (Yang et al., 2006). To gauge if hypoxia affects changes in Itch Tyr-phosphorylation, we immunprecipitated Itch and undertook immunoblotting with phospho-tyrosine (pTyr) specific antibodies. We observed an increase in tyrosine phosphorylation of immunoprecipitated Itch in hypoxia **(Figure 5A)**. To gauge if AMPK is involved in this process, we tested the effects of AMPK knockdown on Itch Tyr phosphorylation. Induction of AMPK knockdown with doxycycline led to a marked decrease in tyrosine phosphorylation of Itch **(Figure 5A)**, revealing a novel role for AMPK in regulating Itch Tyr-phosphorylation in hypoxia.

**Figure 5:**
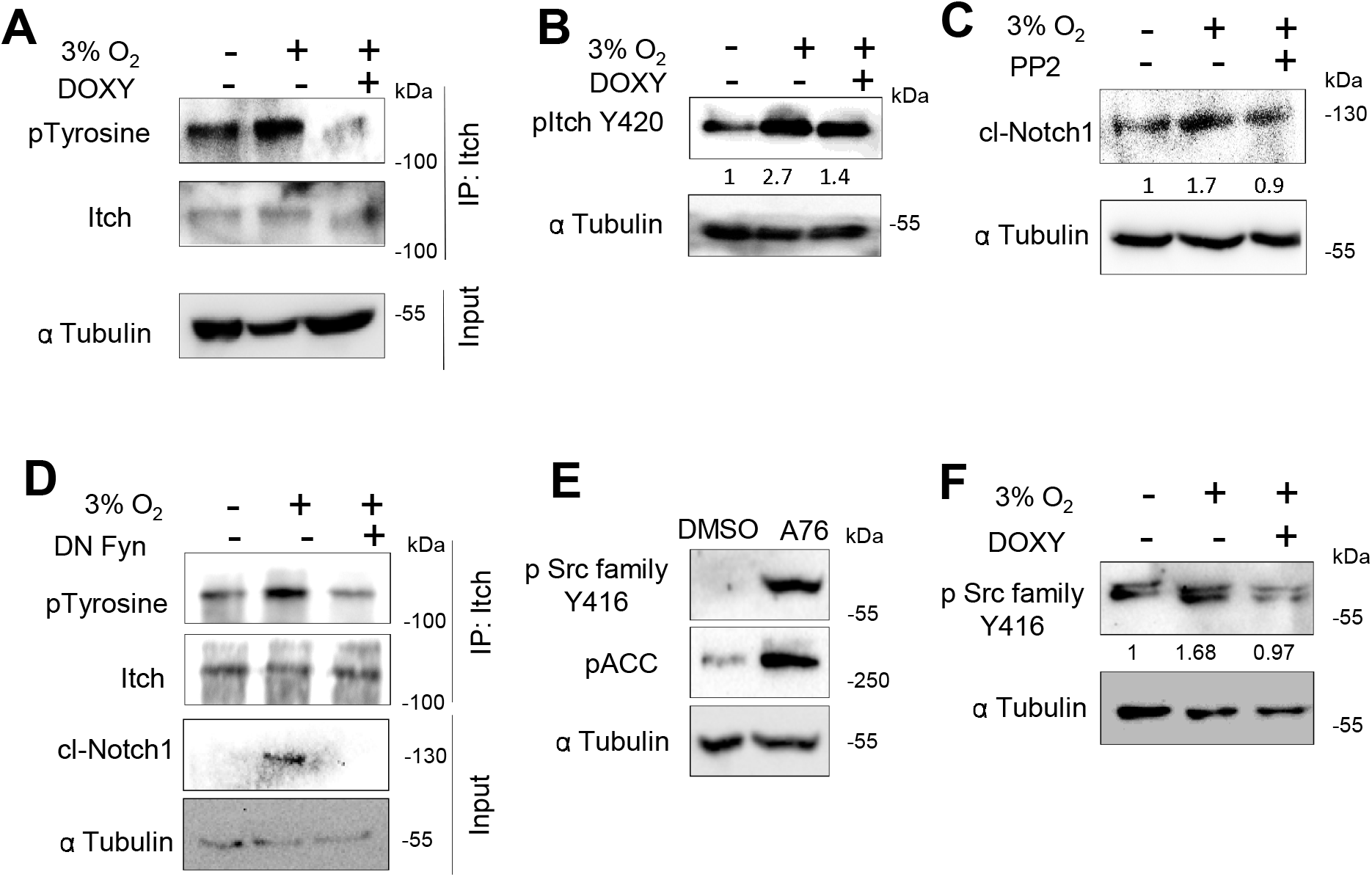
AMPK mediates Itch phosphorylation in hypoxia. (A) MDA-MB-231 cells stably expressing shAMPK α2 (#1) were grown for 24 h in 21% O_2_, or in 3% O_2_ with and without induction by doxycycline (5μg/μl) and Itch was immunoprecipitated and immunoblotted for phospho-tyrosine levels (n=3) (B) MDA-MB-231 cells stably expressing shAMPK α2 (#1) were grown in 21% O_2_ or 3% O_2_ for 24 h with and without induction by doxycycline (5μg/μl) and were immunoblotted for phospho-Itch Y420 levels (this blot was performed by reprobing the blot in Figure 2F, and hence the tubulin panel has been recalled from Figure 2F. (C) MDA-MB-231 cells grown in 21% O_2_ or 3% O_2_ and were treated with PP2 (10 μM) for 24 h and harvested for immunoblotting of specified proteins. (D) MDA-MB-231 cells transfected with control vector or dominant negative Fyn (DN Fyn) were grown in 21% O_2_ or 3% O_2_ for 24 h and harvested for Itch immunoprecipitation followed by immunoblotting for phospho-tyrosine levels (n=3). (E) MDA-MB-231 cells were treated with A76 (100 μM) or DMSO control for 24 h and harvested for immunoblotting and probed for phospho Src family kinase levels (pSFK) (n=3). (F) MDA-MB-231 cells stably expressing shRNA against AMPK α2 (#1) with and without induction by doxycycline (5μg/μl) for 24 h were grown in 3% O_2_ for 24 h harvested for immunoblotting for pSFK levels (n=3).

Since AMPK is a Ser/Thr kinase, whereas we noticed a change in Tyr phosphorylation of Itch in response to AMPK activation, we hypothesized that AMPK might be mediating its effects indirectly, by affecting a tyrosine kinase. Fyn is a Src family tyrosine kinase which is known to phosphorylate and inhibit Itch activity (Yang et al., 2006). To check if Fyn-mediated Itch phosphorylation is involved downstream of AMPK in cleaved Notch1 stabilization in hypoxia, we used an antibody that specifically recognizes tyrosine phosphorylation of human Itch (Tyr 420). We observed an increase in the levels of Itch Y420 phosphorylation under hypoxia **(Figure 5B)**; AMPK knockdown with doxycycline decreased this phosphorylation **(Figure 5B)**. These results identify a novel role for AMPK in Fyn-mediated phosphorylation of Itch in hypoxia.

To further confirm a role for Fyn in AMPK-mediated cleaved Notch1 stabilization in hypoxia we tested the effects of Fyn inhibition on cleaved Notch 1 levels in hypoxia. Treatment of cells with PP2, a Src family kinase inhibitor, hindered the hypoxia-triggered elevation in cleaved Notch1 levels **(Figure 5C)**. To more specifically gauge the role of Fyn, we transfected cells with a dominant negative construct of Fyn and investigated its effect on the tyrosine phosphorylation of Itch in hypoxia. To do so, we immunoprecipitated endogenous Itch and probed for Tyr-phosphorylation by immunoblotting. As shown before, hypoxia led to an increase in Tyr-phosphorylation of Itch, but transfection of DN Fyn impaired this **(Figure 5D)**. Consistent with this, we noticed reduced cleaved Notch1 levels in DN Fyn expressing cells in hypoxia (**Figure 5D**, input WB).

To further confirm the role of Fyn in the regulation of active, cleaved Notch1 in hypoxia, we depleted Fyn using shRNA and tested its effect on NICD levels. Depletion of Fyn led to a reduction in NICD levels in hypoxia **(Figure S5A)**. Together, these results confirmed the involvement of Fyn in stabilizing cleaved Notch1 in hypoxia.

The observations so far revealed an AMPK-dependent and Fyn-mediated negative regulation of Itch in hypoxia. However, a link between AMPK and Fyn activity remains unknown. To address this, we first investigated the effects of AMPK activation and inhibition on Fyn activity. The activity of Src-family kinases (SFK), of which Fyn is a member (Parsons and Parsons, 2004), is regulated by phosphorylation at specific tyrosine residues. Since phospho-Fyn specific antibodies are not available, we used SFK (Tyr 416) recognizing antibody which detects Tyrosine 416 specific phosphorylation on Src-family members that leads to their activation (Roskoski Jr, 2015). AMPK activation with A769662 led to an increase in the levels of tyrosine 416 phosphorylation on Src family kinases **(Figure 5E)**. To understand the significance of this AMPK-mediated positive regulation of Fyn activity in hypoxia, we gauged the effect of AMPK depletion on Src-family tyrosine 416 phosphorylation levels in hypoxia by immunoblotting. In MDA MB 231 cell harbouring the doxycycline-inducible AMPK KD system, in the absence of doxycycline, hypoxia increased phosphorylated Src-family tyrosine 416 levels **(Figure 5F)**. However, addition of doxycycline brought about a reduction in phosphorylated Src-family tyrosine 416 levels **(Figure 5F)**. These data suggested a possible role for AMPK in positively regulating the activity of Fyn in hypoxia.

To further understand the mechanism by which AMPK augments Fyn activation, we carried mass spectrometry to detect interacting proteins of Fyn in cells treated with AMPK activator A769662 and vehicle control DMSO. HEK293T cells stably expressing FLAG-tagged Fyn were immunoprecipitated with FLAG specific antibodies and protein complexes were isolated and subjected to LC-MS. We specifically focussed on the association of Fyn with phosphatases that are known to dephosphorylate Fyn and affect its activity. PTPRF (Vacaresse et al., 2008), a phosphatase known to dephosphorylate the inhibitory phosphorylation at Y528 and thus activate Fyn, was associated with Fyn only in AMPK activated condition, but not in control DMSO treated cells **(Supplementary Table 1)**. Further, PTPN5 (Nguyen et al., 2002) and PTPRC (Vacaresse et al., 2008) are phosphatases known to dephosphorylate the activating phosphorylation of Fyn at Y416, thus decreasing its activity. We detected these in DMSO condition, but not in AMPK activated cells **(Supplementary Table 2)**. Thus, our proteomic analysis further substantiated the role of AMPK in positively modulating Fyn activity by facilitating differential association of phosphatases that promote activation or inactivation of Fyn. These results together indicate an important role for Fyn kinase in AMPK-dependent regulation of Itch activity in hypoxia. Together, these data identify a novel AMPK-dependent regulation of tyrosine phosphorylation of Itch by Fyn kinase in the stabilization of cleaved Notch1.

### Hypoxia-induced AMPK activation promotes stem-like properties and drug-resistance in breast cancer cells through Notch signaling

Having established an interplay between AMPK and Notch, we next investigated the biological significance of the AMPK-Notch pathway interaction in hypoxia. Hypoxic condition is known to facilitate cancer stem cell (CSC)-like properties through involvement of Notch signaling (Sahlgren et al., 2008; Xing et al., 2011). We investigated if AMPK is involved in this process. To address this, we first gauged the effect of AMPK inhibition or depletion on the hypoxia-induced CSC phenotype by measuring the expression of stemness markers like Bmi1 and Nanog. As expected, MDA-MB-231 cells grown in 3% oxygen showed elevated levels of Bmi1 and Nanog compared to normoxia **(Figure 6A)**. However, inhibition of AMPK with compound C in the same experiment prevented the hypoxia-induced elevated expression of these stemness markers **(Figure 6A)**. Likewise, CoCl_2_-induced expression of Bmi1 in these cells was also abrogated by AMPK inhibition **(Figure S6A)**. We also observed similar effect in yet another invasive cancer cell line BT474 **(Figure S6B)**, suggesting a role for AMPK in the hypoxic response leading to elevated stemness.

**Figure 6:**
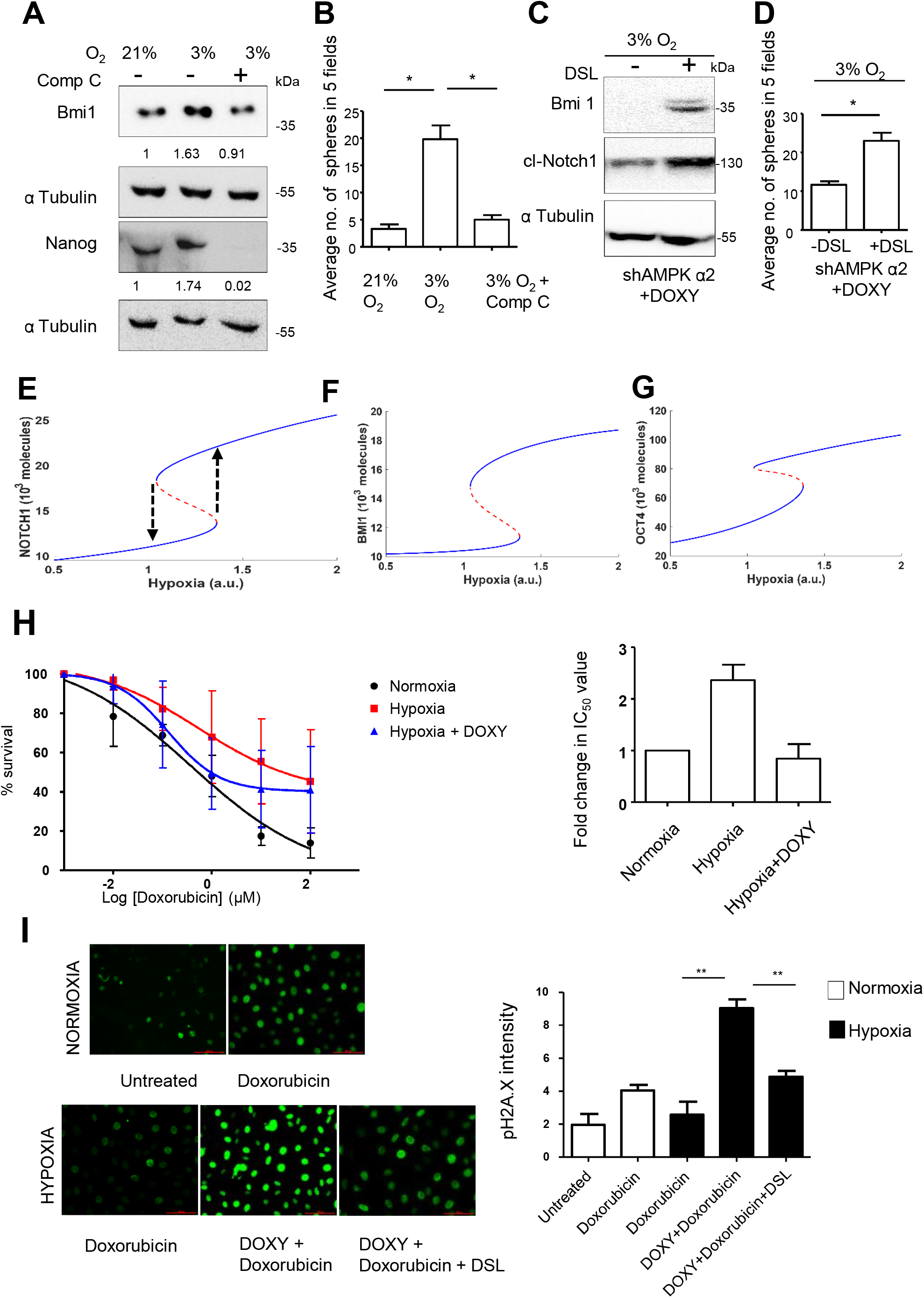
Hypoxia-induced AMPK activation promotes stem-like properties and drug-resistance in breast cancer cells through Notch signaling. (A-B) MDA-MB-231 cells grown in 21% O_2_ or 3% O_2_ were treated with DMSO or Comp C (10 μM) for 48 h and (A) immunoblotted for indicated proteins and (B) subjected to sphere formation assay. The plot represents the average number of day-7 spheres in 5 fields in two replicates of 3 independent experiments. (C-D) MDA-MB-231 cells stably expressing shAMPK α2 (#1) were grown in 3% O_2_ for 24 h with doxycycline induction and subsequently treated with and without DSL ligand (100 nM) and (C) immunoblotted for indicated proteins and (D) subjected to sphere formation assay. The plot represents the average number of day-7 spheres of 3 independent experiments. (E) Bifurcation diagram showing the switch (arrows) from a differentiated state (lower blue curve) to a more stem-like state (upper blue curve) upon increase in hypoxia. (F-G) Bifurcation diagrams for BMI1 and OCT4 showing transitions from a more differentiated state (lower blue curve) to a more stem-like one (upper blue curve) (H) MDA-MB-231 cells stably expressing inducible shAMPK α2 were treated with indicated concentrations of doxorubicin for 48 h under normoxic and hypoxic conditions with and without with doxycycline induced AMPK α2 knockdown (n=3). Cell viability evaluated by MTT assay and fold change in IC_50_ values have been indicated. (I) MDA-MB-231 cells stably expressing inducible shAMPK α2 were treated with 1μM of doxorubicin for 24 h under normoxic and hypoxic conditions with and without with doxycycline induced AMPK α2 knockdown and additional Notch activation by DSL ligand. DNA damage was assessed by pH2A.X staining. Graphs represent the mean intensity of pH2A.X staining from 3 experiments.

To further confirm a role for AMPK in hypoxia-triggered stemness properties, we investigated the effect on sphere formation in defined media condition – an in vitro surrogate for stemness. As expected, hypoxia led to an increase in the number of spheres generated; however, AMPK inhibition blocked this **(Figure 6B)**. We obtained similar results with CoCl_2_ treatment in the presence of AMPK inhibitor **(Figure S6C)**, together indicating the requirement of AMPK in hypoxia-induced sphere forming potential.

Since AMPK depletion downregulated Notch signaling, we asked if increasing cleaved Notch1 levels can rescue the effects of AMPK depletion on stemness properties under hypoxia. To do so, we treated doxycycline induced AMPK depleted MDA-MB-231 cells with a synthetic short peptide (referred to as DSL peptide) corresponding to the Notch ligand Jagged1 whose overexpression promotes Notch signaling (Nickoloff et al., 2002). DSL peptide treatment led to an increase in the levels of cleaved Notch1 in the doxycycline-induced AMPK depleted cells **(Figure 6C)**. Treatment with DSL also led to an increase in the stemness marker Bmi1 expression **(Figure 6C)**, as well as increased the number of spheres formed in these cells **(Figure 6D)**. Integrating our observations with existing literature about stemness regulators, we devised a mathematical model that predicted that hypoxia triggered AMPK can drive as well as maintain a stem-like state characterized by high levels of Notch1 and stemness markers such as Oct4 and Bmi1 (**Figure 6E-G**, **Figure S6D**, Supplementary methods). Put together, these data suggest the involvement of AMPK-Notch signaling axis in the hypoxia-mediated increase in stemness.

An increase in stemness property is also associated with drug resistance (Prieto-Vila et al., 2017), and Notch signaling has been implicated in drug resistance under hypoxic condition in various tumors such as osteosarcoma, ovarian cancer and breast cancer (Morata-Tarifa et al., 2016; Sansone et al., 2007; Seo et al., 2016). We investigated whether the AMPK-Notch1 signaling axis affected drug sensitivity of breast cancer cells under hypoxia. To do so, we first gauged the effect of treatment of doxycycline-inducible AMPK knockdown cells with chemotherapeutic drug doxorubicin under hypoxia. We observed a 2-fold increase in the IC_50_ value for doxorubicin when the cells were exposed to hypoxia in the absence of doxycycline; however, addition of doxycycline prevented this response **(Figure 6H)**.

To further confirm a role for AMPK in hypoxia-induced drug resistance phenotype, we measured the extent of DNA damage accumulated by these cells in hypoxic condition by undertaking pH2A.X staining. As expected, we observed an increase in pH2A.X staining intensity upon treatment with doxorubicin (1μM) under normoxic condition **(Figure 6I)**. In contrast, in hypoxia, treatment with the same concentration of doxorubicin failed to induce DNA damage response **(Figure 6I)**, whereas AMPK-depletion restored the DNA damage response in hypoxia **(Figure 6I)**, suggesting that hypoxia-induced AMPK contributes to drug resistance. To gauge the involvement of Notch in this process, we treated AMPK knockdown cells with synthetic DSL peptide which increases cleaved Notch1 levels in these cells under hypoxia **(Figure 6C)** and assessed the DNA damage. We observed a reduction in pH2A.X staining in DSL peptide-treated cells compared to control untreated cells, indicating restoration of drug resistance phenotype in AMPK depleted cells by increasing cleaved Notch1 levels **(Figure 6I)**. Thus, AMPK-Notch axis plays a key role in mediating CSC phenotype of stemness and drug resistance driven by hypoxia.

### Hypoxia-induced AMPK-Notch1 signaling in vivo

Since hypoxia conditions are prevalent in tumors (Gruber et al., 2004), we next sought to understand the relevance of the AMPK-Notch1 axis in vivo in tumors derived from animal xenografts. To do so, we measured the levels of cleaved and active forms of Notch1 (NICD) using Valine 1744 specific antibody in tumors derived from control and AMPK depleted cells. We used tumors derived from BT 474 cells stably expressing control scrambled constructs or shRNA constructs against AMPK α2 that were injected sub-cutaneously into immunocompromised mice (Hindupur et al., 2014). In small tumors generated by AMPK-depleted cells, we found that NICD levels were highly diminished compared to control scrambled tumors **(Figure 7A)**.

**Figure 7:**
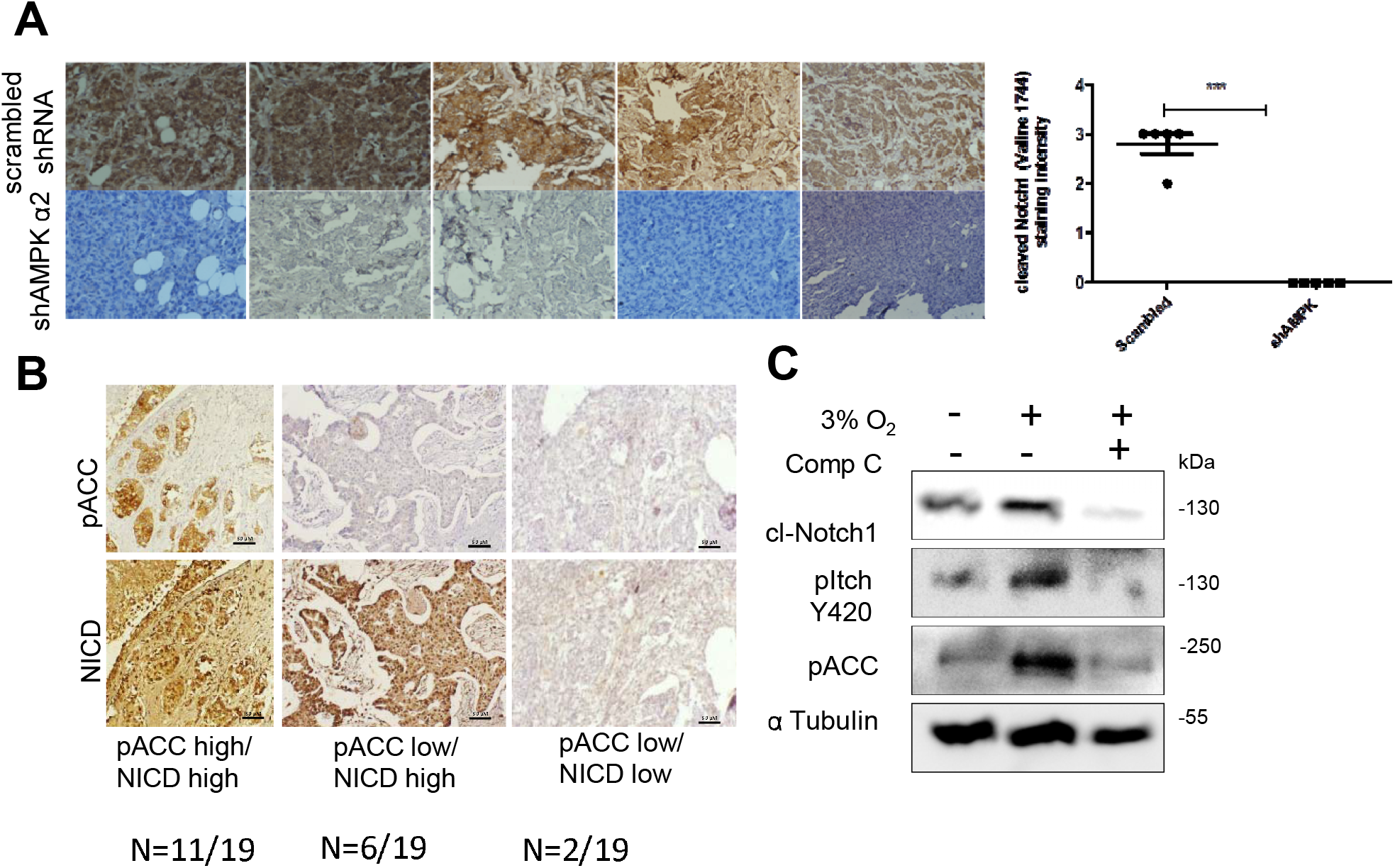
Hypoxia-induced AMPK-Notch1 signaling in vivo. (A) Representative images from immunohistochemical analysis performed on xenograft tumors generated using BT-474 cells stably expressing scrambled or shAMPKα2 to assess cleaved Notch1 (Valine 1744) NICD levels (Scale bar: 100 μm). Graphs represent the staining intensity of cleaved Notch1 (Valine 1744) NICD. (B) Representative images from immunohistochemical analysis of pACC and cleaved Notch1 (Valine 1744) NICD in breast cancer patient samples (n=19). (C) Patient derived breast cancer epithelial cells grown in 21% or 3% O_2_ were treated with DMSO or Comp C (10 μM) for 24 h and immunoblotted for indicated proteins (n=3).

We next addressed the relevance of AMPK-Notch signaling in human breast cancer. High grade breast cancers are majorly associated with hypoxia (Vaupel et al., 2002). We sought to gauge AMPK activity and NICD levels in high grade breast cancer tissue samples. To do so, we performed immunohistochemical analyses using antibodies against pACC (as a measure of AMPK activity) and against gamma secretase cleavage-specific Notch1 (Valine 1744) antibodies (as a measure of NICD) levels. We performed co-staining for pACC and NICD and immunofluorescence microscopy revealed high co-expression in patient samples **(Figure S7A)**. Further, we observed a predominant cytoplasmic staining for pACC while NICD staining was nuclear, as expected **(Figure S7A)**. We then performed DAB-based immunohistochemistry in serial sections of chemo-naïve, grade III invasive ductal carcinoma breast cancer patient sample (N=19) for these two antibodies. We observed an association (p=0.0422, Fischer exact test) between pACC and NICD levels in these samples **(Figure 7B)**. Together, these data highlighted the importance of the AMPK-Notch1 axis in patient tumors.

Furthermore, we investigated into the response of patient-derived breast cancer cells to hypoxia. The primary tissue-derived cells exposed to hypoxic condition showed elevated levels of pACC, cleaved Notch1 and phospho-Itch levels, whereas pharmacological inhibition of AMPK in these cells under hypoxic condition reduced the cleaved Notch1 and phospho-Itch levels **(Figure 7C)**, suggesting that the AMPK-Itch-Notch axis might also operate in patient-derived cells.

In order to further understand the clinical relevance of our study, we analysed the gene signatures of AMPK (22), Hypoxia (40) and Notch (26) pathways (Supplementary Table 3) in TCGA primary breast cancer dataset by performing gene-wise correlation across the signatures by Pearson’s product moment correlation **(Figure 8 A-C)** and Spearman correlation methods **(Figure S8)**. We observed significant correlation (Pearson co-relation) between AMPK-Hypoxia, Notch-Hypoxia and AMPK-Notch gene signatures **(Figure 8A-C)**. There was no difference observed between both the correlation methods used (Pearson vs Spearman’s co-relation) **(Figures 8 & S8)**. In 880 (22*40), 1040 (26*40), 572 (22*26) number of pair-wise gene correlations between AMPK-Hypoxia **(Figure 8A)**, Notch-Hypoxia **(Figure 8B)**, and AMPK-Notch **(Figure 8C)** respectively, 669, 840 and 515 gene pairs showed a positive correlation, of which the correlation for 534, 669, 412 gene pairs were significant (p<0.01), implying a strong interaction between hypoxia-AMPK-Notch signaling pathways. Further, hierarchical clustering analysis of gene expression profiles from NCBI-GEO dataset (GSE40206), a microarray dataset of 80 Indian breast cancer patient cohort, revealed co-expression of hypoxia, AMPK and Notch-responsive genes in majority of cases **(Figure S9)**, thus underscoring the relevance of the hypoxia-AMPK-Notch axis in breast cancers.

**Figure 8:**
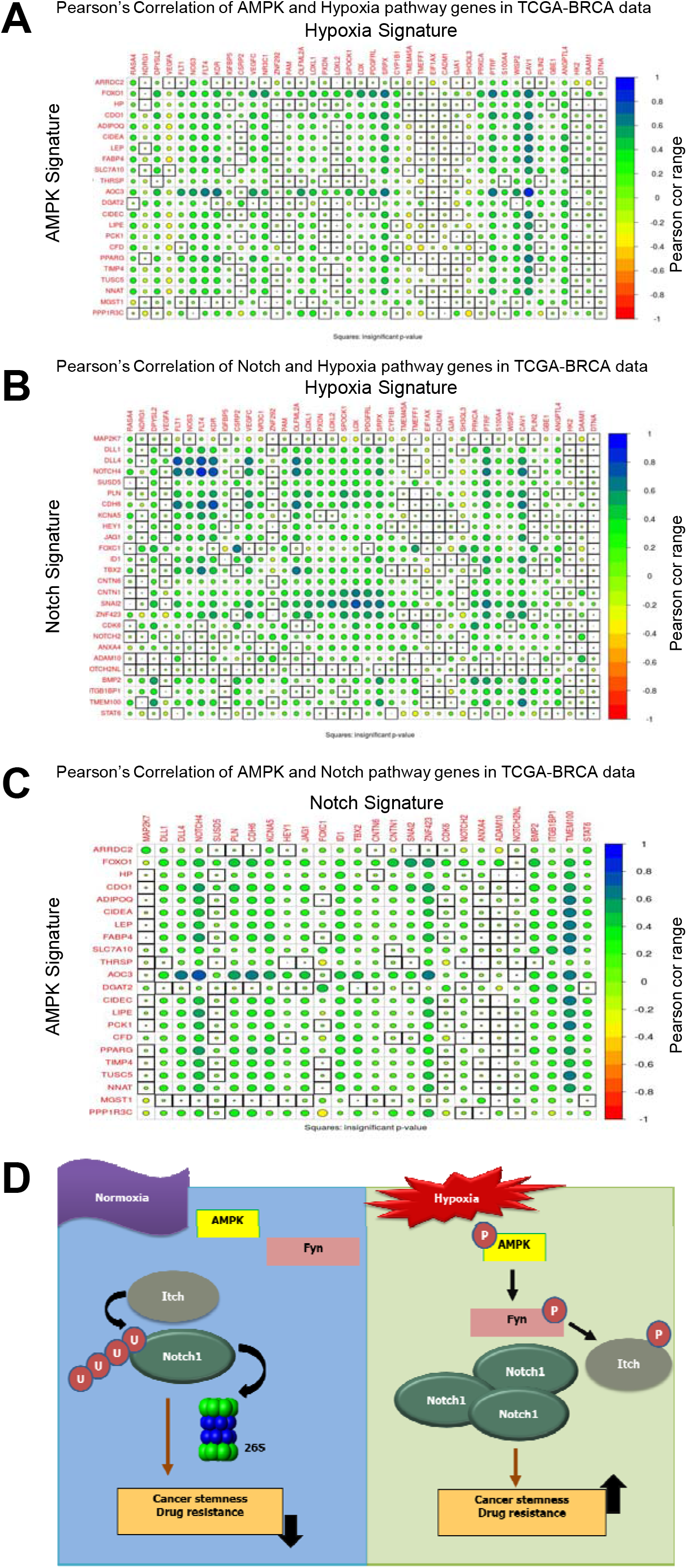
Hypoxia-AMPK-Notch gene signature correlations in TCGA Breast cancer dataset. (A-C) Pearson’s correlation plots between AMPK-Hypoxia (A), Notch-Hypoxia (B) and AMPK-Notch (C) analyzed using AMPK, Notch and Hypoxia gene signature in TCGA-BRCA (all subtypes) dataset; Pearson’s correlation value (cor) of each gene pair is represented as the size of the circle and filled with corresponding color from the color palette represented below the ranging from −1(red) to +1(blue). Boxes highlighted by the black squares represent insignificant (p>0.01) Pearson cor values. (D) Proposed model depicting AMPK and Notch crosstalk under normoxia and hypoxia: In conditions of normoxia, basal activity of AMK does not interfere with interaction of E3-ubiquitin ligase Itch with its substrate cleaved Notch1, which is ubiquitinated and targeted for proteasomal degradation. In conditions of hypoxia, activated AMPK levels enhance Fyn activation, which phosphorylates and hinders binding of ligase Itch with its substrate cleaved Notch1, thereby resulting in cleaved Notch1 stabilization. This results in elevated stemness and drug resistance of the tumors.

## Discussion

Hypoxia in solid tumors is considered a malice aiding cancer progression and hindering successful therapy. Notch signaling is implicated as a major regulator of hypoxia-mediated stemness and drug resistance. Because of its importance in numerous cancers, the Notch pathway has been a major candidate for developing therapeutic agents. While gamma secretase inhibitors and antibodies against receptor and ligands have reached clinical trials, gastrointestinal toxicity and other side effects have led to shifting of focus on druggable targets that impinge on the regulation of Notch signaling (Andersson and Lendahl, 2014; Guo et al., 2011). In this study, we identify AMPK as a novel molecular regulator of cleaved Notch1 stability under hypoxia (Figure 8D), and propose that inhibition of AMPK can be a potential therapeutic strategy impairing the adaptation of breast cancer cells to hypoxia.

Hypoxia-induced Notch1 signaling has been reported in melanoma (Bedogni et al., 2008), adenocarcinoma of lungs (Chen et al., 2007), glioma (Qiang et al., 2012), as well as breast carcinomas (Sahlgren et al., 2008). Mechanistically, it has been shown that HIF1α and NICD directly interact and augment transcription of Notch responsive genes (Gustafsson et al., 2005). Another mechanism implicated in this crosstalk involves FIH (factor inhibiting HIF1-α) which is critical in blocking differentiation in myogenic and neural progenitor cells (Zheng et al., 2008). HIF-α synergises with Notch co-activator MAML to increase expression of *Snail* and *Slug* and reduce *E-cadherin* expression (Chen et al., 2010). Treatment with mTORC1/2 inhibitors also elevate Notch signaling and maintain a drug-resistant CSC phenotype (Bhola et al., 2016) in triple-negative breast cancers. Only one study has reported the stabilization and increased half-life of NICD in hypoxia in a HIF1α-dependent manner (Gustafsson et al., 2005); however, the molecular mechanisms remained unknown. We show here that AMPK activation is critical for cleaved Notch1 stabilization under hypoxia. Using cycloheximide chase assays, we show that AMPK inhibition, depletion or genetic ablation causes enhanced degradation of cleaved Notch1. Cleaved Notch1 is targeted for proteasomal degradation after ubiquitination (Öberg et al., 2001; Qiu et al., 2000; Wu et al., 2001). We show here that the cleaved Notch1-K-48-linked ubiquitination, which marks proteins for degradation, is reduced under hypoxia as well as upon pharmacological AMPK activation. In contrast, AMPK inhibition or depletion increases ubiquitination of cleaved Notch1. We further show that in AMPK DKO MEFs which do not have AMPK activity, cleaved Notch levels are low, which can be restored by AMPK re-constitution. Our study thus identifies a novel role for AMPK in cleaved Notch1 stabilization. Our data revealed that CoCl_2_ treatment led to both AMPK activation as well as elevated cleaved Notch1 levels. Furthermore, AMPK inhibition led to heightened decrease of CoCl_2_-induced elevated cleaved Notch1 levels. Since CoCl_2_ mimics hypoxia by stabilizing HIF-1α, these data suggest a possible role for HIF1-α in this process. However, hypoxia can trigger AMPK both in HIF-1α dependent (Singh et al., 2013) as well independent fashion (Marin et al., 2016). Thus, AMPK and HIF-1α are likely to affect cleaved Notch1 stability independently, as well as in a concerted manner. Further investigations are required to distinguish their individual and combined effects.

Post-translational modifications like phosphorylation and acetylation of the Notch intracellular domain regulate its ubiquitination (Fryer et al., 2004; Popko-Scibor et al., 2011). However, we failed to find significant changes in phosphorylation or acetylation of cleaved Notch upon AMPK activation. Notch and its interaction with ubiquitin ligases has been often shown to be influenced by regulators like Numb, which suppresses Notch signaling by recruiting Itch and facilitating degradation of Notch (McGill and McGlade, 2003). We identified AMPK as a negative regulator of Itch-mediated Notch degradation. Several studies have identified phosphorylation of Itch as a major regulator of its activity. Kinases like ATM and JNK activate Itch by increasing its enzymatic activity for a specific set of targets like c-Jun or c-FLIP, whereas other kinases like Fyn reduce interaction of Itch with substrates including Jun-B as well as Notch leading to lesser ubiquitination (Melino et al., 2008). A previous study showed an inhibitory effect of AMPK activator on interaction of Itch with its substrate p73 leading to lesser degradation of the latter (Adamovich et al., 2014). However, the mechanisms that lead to AMPK-dependent alteration of Itch activity was not explored. We show that AMPK activation does not alter Itch levels, but it impairs Itch-Notch interaction by the regulation of Itch tyrosine phosphorylation by Fyn **(Figure 8D)**. We also show that AMPK promotes Fyn activation by altering its interaction with tyrosine phosphatases that regulate its activity. This is in keeping with a report showing LKB1, an upstream kinase of AMPK, promotes protein tyrosine phosphatase activity leading to inhibition of receptor tyrosine kinases in lung and cervical cancer cell lines (Okon et al., 2014). Likewise, we recently reported AMPK-mediated regulation of serine threonine kinase Akt activity by modulating its phosphatase PHLPP2 in breast cancer cells (Saha et al., 2018). Thus, AMPK might alter the activity of a vast number of other kinases by affecting their interaction with or activity of cellular phosphatases. These observations warrant a detailed investigation into AMPK-mediated regulation of signal transduction pathways by regulating cellular phosphatases.

AMPK is traditionally viewed as a tumor suppressor as it negatively regulates mTOR, thus bringing about slower growth (Gwinn et al., 2008). AMPK is also known to stabilize p53 bringing about cell cycle arrest (Zadra et al., 2015). Both mTOR and p53 pathways have been shown to cross talk with Notch. mTOR has been reported to regulate Notch1 transcription (Bhola et al., 2016), but we failed to see changes in Notch1 transcript levels upon AMPK activation. While p53 was shown to positively regulate Notch1 in epithelial cells (Yugawa et al., 2007), we have tested the hypoxia-AMPK-Notch signaling axis in p53 mutant cell line (MDA-MB-231) indicating that p53 is not involved in mediating the observed effects of AMPK on cleaved Notch1 levels. We show a pro-tumorigenic role of AMPK through positive regulation of Notch1 stability leading to elevated breast cancer stemness and drug resistance phenotype **(Figure 8D)**. Interestingly, our study shows a high correlation between AMPK and Notch activation in grade 3 primary breast cancer samples that are known to contain hypoxic regions (Zhang et al., 2016). In support of our observations, abundant expression of activated AMPK has also been reported in human gliomas (Ríos et al., 2013) and breast cancer (Hart et al., 2015a). Contextual oncogenic properties of AMPK has been reported, especially in bioenergetic stress conditions such as nutrient deprivation and hypoxia, which are often faced by cells of a growing tumor (Kishton et al., 2016; Liang and Mills, 2013). While we have identified a role for AMPK in increasing breast cancer stemness and drug resistance in hypoxia, previously, a protective role for AMPK has been demonstrated in androgen-dependent prostate cancer cells in hypoxia (Chhipa et al., 2011) and in other solid tumors with low oxygen condition (Laderoute et al., 2006). Further, studies from our lab (Hindupur et al., 2014) and others (Jeon et al., 2012) show in-vivo evidence of tumor promoting role of AMPK in tumor xenograft models. Another study showed that this role of AMPK is evident only in a metabolically stressed tumor microenvironment context (Laderoute et al., 2014). Thus, our report compliments such studies on tumor promoting role of AMPK in the context of tumor hypoxia.

We have recently reported the role of AMPK in supporting EMT and metastasis in hypoxia (Saxena et al., 2018). An increased metastatic potential is associated with drug resistance. Consistent with this, recent studies have implicated AMPK in mediating hypoxia-induced drug resistance to doxorubicin in osteosarcoma (Zhao et al., 2016). Other studies have reported AMPK in resistance to cisplatin (Harhaji□Trajkovic et al., 2009; Shin et al., 2014). In this study, we show that hypoxia-induced drug resistance phenotype mediated by Notch signaling in breast cancer cells is supported by AMPK activation.

Our study identifies a novel protective role for AMPK in Itch-mediated Notch degradation. We further show that AMPK plays an important role in maintaining the stemness and drug resistance phenotypes induced by hypoxia. Thus, under the stress of hypoxia which is prevalent in solid tumors, we propose that AMPK inhibition-based strategies can alleviate the cancer stem cell phenotype induced by hypoxia, thereby rendering cancer cells more susceptible to existing anti-cancer agents.

## Acknowledgement

Funding: This work was majorly supported by grants from the Wellcome Trust-DBT India Alliance (IA) Senior Research Fellowship (500112/Z/09/Z) to AR. ML acknowledges Council for Scientific and Industrial Research for CSIR fellowship (19-12/2010(i) EU-IV). MKJ acknowledges support from Ramanujan Fellowship provided by SERB, DST, Govt. of India (SB/S2/RJN-049/2019). The authors acknowledge grants from DBT-IISc partnership programme, support from DST-FIST and UGC, Govt. of India to the Department of MRDG, IISc.

The authors thank Dr. Benoit Viollet for DKO MEFs and Sai Balaji for help with xenograft assay. The authors acknowledge the Central Animal Facility and FACS facility at IISc.

Competing interests: The authors declare that they have no competing interests.

## Authors’ contributions

ML and AR conceived and designed experiments; ML performed majority of the experiments and analyzed data; SK performed some experiments, and helped with bioinformatics data analysis. AC performed correlation analyses from TCGA data set. KH and MKJ devised the mathematical model. ML and AR edited and drafted the manuscript. AR supervised the study. All authors have read and approved the final version of the manuscript.

## Declaration

Ethics approval and consent to participate: Primary breast tissues (cancer and adjacent normal) obtained from Kidwai Memorial Institute of Oncology (KMIO), Bangalore, as per IRB and in compliance with ethical guidelines of KMIO and the Indian Institute of Science (IISc). All animal experiments were reviewed and approved by the Institutional Animal Ethics committee of IISc, Bangalore.

Availability of data and material: All data generated or analysed during this study are included in this published article [and its supplementary information files].

## References

Adamovich, Y., Adler, J., Meltser, V., Reuven, N., and Shaul, Y. (2014). AMPK couples p73 with p53 in cell fate decision. Cell Death & Differentiation 21, 1451–1459.

Allenspach, E.J., Maillard, I., Aster, J.C., and Pear, W.S. (2002). Notch signaling in cancer. Cancer biology & therapy 1, 466–476.

Andersson, E.R., and Lendahl, U. (2014). Therapeutic modulation of Notch signalling—are we there yet? Nature reviews Drug discovery 13, 357–378.

Andersson, E.R., Sandberg, R., and Lendahl, U. (2011). Notch signaling: simplicity in design, versatility in function. Development 138, 3593–3612.

Artavanis-Tsakonas, S., Rand, M.D., and Lake, R.J. (1999). Notch signaling: cell fate control and signal integration in development. Science 284, 770–776.

Aster, J.C., Pear, W.S., and Blacklow, S.C. (2008). Notch signaling in leukemia. Annu Rev Pathol Mech Dis 3, 587–613.

Bedogni, B., Warneke, J.A., Nickoloff, B.J., Giaccia, A.J., and Powell, M.B. (2008). Notch1 is an effector of Akt and hypoxia in melanoma development. The Journal of clinical investigation 118, 3660–3670.

Bhola, N.E., Jansen, V.M., Koch, J.P., Li, H., Formisano, L., Williams, J.A., Grandis, J.R., and Arteaga, C.L. (2016). Treatment of triple-negative breast cancer with TORC1/2 inhibitors sustains a drug-resistant and notch-dependent cancer stem cell population. Cancer research 76, 440–452.

Blokzijl, A., Dahlqvist, C., Reissmann, E., Falk, A., Moliner, A., Lendahl, U., and Ibáñez, C.F. (2003). Cross-talk between the Notch and TGF-β signaling pathways mediated by interaction of the Notch intracellular domain with Smad3. The Journal of cell biology 163, 723–728.

Bray, S.J. (2006). Notch signalling: a simple pathway becomes complex. Nature reviews Molecular cell biology 7, 678–689.

Brou, C., Logeat, F., Gupta, N., Bessia, C., LeBail, O., Doedens, J.R., Cumano, A., Roux, P., Black, R.A., and Israël, A. (2000). A novel proteolytic cleavage involved in Notch signaling: the role of the disintegrin-metalloprotease TACE. Molecular cell 5, 207–216.

Chen, J., Imanaka, N., and Griffin, J. (2010). Hypoxia potentiates Notch signaling in breast cancer leading to decreased E-cadherin expression and increased cell migration and invasion. British Journal of Cancer 102, 351–360.

Chen, R., Jiang, T., She, Y., Xu, J., Li, C., Zhou, S., Shen, H., Shi, H., and Liu, S. (2017). Effects of cobalt chloride, a hypoxia-mimetic agent, on autophagy and atrophy in skeletal C2C12 myotubes. BioMed research international 2017.

Chen, Y., De Marco, M.A., Graziani, I., Gazdar, A.F., Strack, P.R., Miele, L., and Bocchetta, M. (2007). Oxygen concentration determines the biological effects of NOTCH-1 signaling in adenocarcinoma of the lung. Cancer research 67, 7954–7959.

Chhipa, R.R., Wu, Y., and Ip, C. (2011). AMPK-mediated autophagy is a survival mechanism in androgen-dependent prostate cancer cells subjected to androgen deprivation and hypoxia. Cellular signalling 23, 1466–1472.

Cool, B., Zinker, B., Chiou, W., Kifle, L., Cao, N., Perham, M., Dickinson, R., Adler, A., Gagne, G., and Iyengar, R. (2006). Identification and characterization of a small molecule AMPK activator that treats key components of type 2 diabetes and the metabolic syndrome. Cell metabolism 3, 403–416.

D’Angelo, R.C., Ouzounova, M., Davis, A., Choi, D., Tchuenkam, S.M., Kim, G., Luther, T., Quraishi, A.A., Senbabaoglu, Y., and Conley, S.J. (2015). Notch reporter activity in breast cancer cell lines identifies a subset of cells with stem cell activity. Molecular cancer therapeutics 14, 779–787.

Elenbaas, B., Spirio, L., Koerner, F., Fleming, M.D., Zimonjic, D.B., Donaher, J.L., Popescu, N.C., Hahn, W.C., and Weinberg, R.A. (2001). Human breast cancer cells generated by oncogenic transformation of primary mammary epithelial cells. Genes & development 15, 50–65.

Emerling, B.M., Weinberg, F., Snyder, C., Burgess, Z., Mutlu, G.M., Viollet, B., Budinger, G.S., and Chandel, N.S. (2009). Hypoxic activation of AMPK is dependent on mitochondrial ROS but independent of an increase in AMP/ATP ratio. Free Radical Biology and Medicine 46, 1386–1391.

Fryer, C.J., White, J.B., and Jones, K.A. (2004). Mastermind recruits CycC: CDK8 to phosphorylate the Notch ICD and coordinate activation with turnover. Molecular cell 16, 509–520.

Gordon, W.R., Arnett, K.L., and Blacklow, S.C. (2008). The molecular logic of Notch signaling–a structural and biochemical perspective. Journal of cell science 121, 3109–3119.

Gruber, G., Greiner, R.H., Hlushchuk, R., Aebersold, D.M., Altermatt, H.J., Berclaz, G., and Djonov, V. (2004). Hypoxia-inducible factor 1 alpha in high-risk breast cancer: an independent prognostic parameter? Breast Cancer Research 6, 1.

Guo, S., Liu, M., and Gonzalez-Perez, R.R. (2011). Role of Notch and its oncogenic signaling crosstalk in breast cancer. Biochimica et Biophysica Acta (BBA)-Reviews on Cancer 1815, 197–213.

Gupta-Rossi, N., Le Bail, O., Brou, C., Logeat, F., Six, E., and Israël, A. (2002). Control of Notch Activity by the Ubiquitin-Proteasome Pathway. In Notch from Neurodevelopment to Neurodegeneration: Keeping the Fate (Springer), pp. 41–58.

Gusarova, G.A., Trejo, H.E., Dada, L.A., Briva, A., Welch, L.C., Hamanaka, R.B., Mutlu, G.M., Chandel, N.S., Prakriya, M., and Sznajder, J.I. (2011). Hypoxia leads to Na, K-ATPase downregulation via Ca2+ release-activated Ca2+ channels and AMPK activation. Molecular and cellular biology 31, 3546–3556.

Gustafsson, M.V., Zheng, X., Pereira, T., Gradin, K., Jin, S., Lundkvist, J., Ruas, J.L., Poellinger, L., Lendahl, U., and Bondesson, M. (2005). Hypoxia requires notch signaling to maintain the undifferentiated cell state. Developmental cell 9, 617–628.

Gwinn, D.M., Shackelford, D.B., Egan, D.F., Mihaylova, M.M., Mery, A., Vasquez, D.S., Turk, B.E., and Shaw, R.J. (2008). AMPK phosphorylation of raptor mediates a metabolic checkpoint. Molecular cell 30, 214–226.

Hamilton, S.R., Stapleton, D., O’Donnell, J.B., Kung, J.T., Dalal, S.R., Kemp, B.E., and Witters, L.A. (2001). An activating mutation in the γ1 subunit of the AMP-activated protein kinase. FEBS letters 500, 163–168.

Hardie, D.G. (2013). AMPK: a target for drugs and natural products with effects on both diabetes and cancer. Diabetes 62, 2164–2172.

Hardie, D.G., and Alessi, D.R. (2013). LKB1 and AMPK and the cancer-metabolism link-ten years after. BMC biology 11, 36.

Hardie, D.G., Ross, F.A., and Hawley, S.A. (2012). AMPK: a nutrient and energy sensor that maintains energy homeostasis. Nature reviews Molecular cell biology 13, 251–262.

Harhaji-Trajkovic, L., Vilimanovich, U., Kravic-Stevovic, T., Bumbasirevic, V., and Trajkovic, V. (2009). AMPK-mediated autophagy inhibits apoptosis in cisplatin-treated tumour cells. Journal of cellular and molecular medicine 13, 3644–3654.

Hart, P., Mao, M., Ansenberger-Fricano, K., Ekoue, D., Ganini, D., Kajdacsy-Balla, A., Diamond, A., Minshall, R., Consolaro, M., and Santos, J. (2015a). MnSOD upregulation sustains the Warburg effect via mitochondrial ROS and AMPK-dependent signalling in cancer. Nature communications 6, 6053–6053.

Hart, P.C., Mao, M., De Abreu, A.L.P., Ansenberger-Fricano, K., Ekoue, D.N., Ganini, D., Kajdacsy-Balla, A., Diamond, A.M., Minshall, R.D., and Consolaro, M.E. (2015b). MnSOD upregulation sustains the Warburg effect via mitochondrial ROS and AMPK-dependent signalling in cancer. Nature communications 6, 6053.

Hindupur, S.K., Balaji, S.A., Saxena, M., Pandey, S., Sravan, G.S., Heda, N., Kumar, M.V., Mukherjee, G., Dey, D., and Rangarajan, A. (2014). Identification of a novel AMPK-PEA15 axis in the anoikis-resistant growth of mammary cells. Breast Cancer Research 16, 1.

Jarriault, S., Brou, C., Logeat, F., Schroeter, E.H., Kopan, R., and Israel, A. (1995). Signalling downstream of activated mammalian Notch. Nature 377, 355–358.

Jeon, S.-M., Chandel, N.S., and Hay, N. (2012). AMPK regulates NADPH homeostasis to promote tumour cell survival during energy stress. Nature 485, 661–665.

Jeon, S.-M., and Hay, N. (2012). The dark face of AMPK as an essential tumor promoter. Cellular logistics 2, 197–202.

Jeon, S.-M., and Hay, N. (2015). The double-edged sword of AMPK signaling in cancer and its therapeutic implications. Archives of pharmacal research 38, 346–357.

Jubb, A., Turley, H., Moeller, H., Steers, G., Han, C., Li, J., Leek, R., Tan, E., Singh, B., and Mortensen, N. (2009). Expression of delta-like ligand 4 (Dll4) and markers of hypoxia in colon cancer. British journal of cancer 101, 1749–1757.

Kim, J.-Y., and Lee, J.-Y. (2017). Targeting tumor adaption to chronic hypoxia: implications for drug resistance, and how it can be overcome. International journal of molecular sciences 18, 1854.

Kishton, R.J., Barnes, C.E., Nichols, A.G., Cohen, S., Gerriets, V.A., Siska, P.J., Macintyre, A.N., Goraksha-Hicks, P., De Cubas, A.A., and Liu, T. (2016). AMPK is essential to balance glycolysis and mitochondrial metabolism to control T-ALL cell stress and survival. Cell metabolism 23, 649–662.

Kunz, M., and Ibrahim, S.M. (2003). Molecular responses to hypoxia in tumor cells. Molecular cancer 2, 23.

Laderoute, K.R., Amin, K., Calaoagan, J.M., Knapp, M., Le, T., Orduna, J., Foretz, M., and Viollet, B. (2006). 5⍰-AMP-activated protein kinase (AMPK) is induced by low-oxygen and glucose deprivation conditions found in solid-tumor microenvironments. Molecular and cellular biology 26, 5336–5347.

Laderoute, K.R., Calaoagan, J.M., Chao, W.-r., Dinh, D., Denko, N., Duellman, S., Kalra, J., Liu, X., Papandreou, I., and Sambucetti, L. (2014). 5⍰-AMP-activated protein kinase (AMPK) supports the growth of aggressive experimental human breast cancer tumors. Journal of Biological Chemistry 289, 22850–22864.

Liang, J., and Mills, G.B. (2013). AMPK: a contextual oncogene or tumor suppressor? Cancer research 73, 2929–2935.

Marin, J.J., Lozano, E., and Perez, M.J. (2016). Lack of mitochondrial DNA impairs chemical hypoxia-induced autophagy in liver tumor cells through ROS-AMPK-ULK1 signaling dysregulation independently of HIF-1α. Free Radical Biology and Medicine 101, 71–84.

McGill, M.A., and McGlade, C.J. (2003). Mammalian numb proteins promote Notch1 receptor ubiquitination and degradation of the Notch1 intracellular domain. Journal of Biological Chemistry 278, 23196–23203.

Melino, G., Gallagher, E., Aqeilan, R., Knight, R., Peschiaroli, A., Rossi, M., Scialpi, F., Malatesta, M., Zocchi, L., and Browne, G. (2008). Itch: a HECT-type E3 ligase regulating immunity, skin and cancer. Cell Death & Differentiation 15, 1103–1112.

Mihaylova, M.M., and Shaw, R.J. (2011). The AMPK signalling pathway coordinates cell growth, autophagy and metabolism. Nature cell biology 13, 1016–1023.

Mittal, S., Subramanyam, D., Dey, D., Kumar, R.V., and Rangarajan, A. (2009). Cooperation of Notch and Ras/MAPK signaling pathways in human breast carcinogenesis. Molecular cancer 8, 1.

Morata-Tarifa, C., Jiménez, G., García, M.A., Entrena, J.M., Griñán-Lisón, C., Aguilera, M., Picon-Ruiz, M., and Marchal, J.A. (2016). Low adherent cancer cell subpopulations are enriched in tumorigenic and metastatic epithelial-to-mesenchymal transition-induced cancer stem-like cells. Scientific reports 6, 18772.

Moretti, J., and Brou, C. (2013). Ubiquitinations in the notch signaling pathway. International journal of molecular sciences 14, 6359–6381.

Mungai, P.T., Waypa, G.B., Jairaman, A., Prakriya, M., Dokic, D., Ball, M.K., and Schumacker, P.T. (2011). Hypoxia triggers AMPK activation through reactive oxygen species-mediated activation of calcium release-activated calcium channels. Molecular and cellular biology 31, 3531–3545.

Nguyen, T.-H., Liu, J., and Lombroso, P.J. (2002). Striatal enriched phosphatase 61 dephosphorylates Fyn at phosphotyrosine 420. Journal of Biological Chemistry 277, 24274–24279.

Nickoloff, B., Qin, J., Chaturvedi, V., Denning, M., Bonish, B., and Miele, L. (2002). Jagged-1 mediated activation of notch signaling induces complete maturation of human keratinocytes through NF-κB and PPARγ. Cell Death & Differentiation 9, 842–855.

Öberg, C., Li, J., Pauley, A., Wolf, E., Gurney, M., and Lendahl, U. (2001). The Notch intracellular domain is ubiquitinated and negatively regulated by the mammalian Sel-10 homolog. Journal of Biological Chemistry 276, 35847–35853.

Okon, I.S., Coughlan, K.A., and Zou, M.-H. (2014). Liver kinase B1 expression promotes phosphatase activity and abrogation of receptor tyrosine kinase phosphorylation in human cancer cells. Journal of Biological Chemistry 289, 1639–1648.

Parsons, S.J., and Parsons, J.T. (2004). Src family kinases, key regulators of signal transduction. Oncogene 23, 7906.

Pece, S., Serresi, M., Santolini, E., Capra, M., Hulleman, E., Galimberti, V., Zurrida, S., Maisonneuve, P., Viale, G., and Di Fiore, P.P. (2004). Loss of negative regulation by Numb over Notch is relevant to human breast carcinogenesis. The Journal of cell biology 167, 215–221.

Popko-Scibor, A.E., Lindberg, M.J., Hansson, M.L., Holmlund, T., and Wallberg, A.E. (2011). Ubiquitination of Notch1 is regulated by MAML1-mediated p300 acetylation of Notch1. Biochemical and biophysical research communications 416, 300–306.

Prieto-Vila, M., Takahashi, R.-u., Usuba, W., Kohama, I., and Ochiya, T. (2017). Drug resistance driven by cancer stem cells and their niche. International journal of molecular sciences 18, 2574.

Qiang, L., Wu, T., Zhang, H., Lu, N., Hu, R., Wang, Y., Zhao, L., Chen, F., Wang, X., and You, Q. (2012). HIF-1α is critical for hypoxia-mediated maintenance of glioblastoma stem cells by activating Notch signaling pathway. Cell Death & Differentiation 19, 284–294.

Qiu, L., Joazeiro, C., Fang, N., Wang, H.-Y., Elly, C., Altman, Y., Fang, D., Hunter, T., and Liu, Y.-C. (2000). Recognition and ubiquitination of Notch by Itch, a hect-type E3 ubiquitin ligase. Journal of Biological Chemistry 275, 35734–35737.

Radtke, F., and Raj, K. (2003). The role of Notch in tumorigenesis: oncogene or tumour suppressor? Nat Rev Cancer 3, 756–767.

Reedijk, M., Odorcic, S., Chang, L., Zhang, H., Miller, N., McCready, D.R., Lockwood, G., and Egan, S.E. (2005). High-level coexpression of JAG1 and NOTCH1 is observed in human breast cancer and is associated with poor overall survival. Cancer research 65, 8530–8537.

Ríos, M., Foretz, M., Viollet, B., Prieto, A., Fraga, M., Costoya, J.A., and Señarís, R. (2013). AMPK activation by oncogenesis is required to maintain cancer cell proliferation in astrocytic tumors. Cancer research 73, 2628–2638.

Roskoski Jr, R. (2015). Src protein-tyrosine kinase structure, mechanism, and small molecule inhibitors. Pharmacological research 94, 9–25.

Saha, M., Kumar, S., Bukhari, S., Balaji, S.A., Kumar, P., Hindupur, S.K., and Rangarajan, A. (2018). AMPK–Akt double-negative feedback loop in breast cancer cells regulates their adaptation to matrix deprivation. Cancer research 78, 1497–1510.

Sahlgren, C., Gustafsson, M.V., Jin, S., Poellinger, L., and Lendahl, U. (2008). Notch signaling mediates hypoxia-induced tumor cell migration and invasion. Proceedings of the National Academy of Sciences 105, 6392–6397.

Samanta, D., Park, Y., Andrabi, S.A., Shelton, L.M., Gilkes, D.M., and Semenza, G.L. (2016). PHGDH expression is required for mitochondrial redox homeostasis, breast cancer stem cell maintenance, and lung metastasis. Cancer research 76, 4430–4442.

Sansone, P., Storci, G., Giovannini, C., Pandolfi, S., Pianetti, S., Taffurelli, M., Santini, D., Ceccarelli, C., Chieco, P., and Bonafe, M. (2007). p66Shc/Notch-3 Interplay Controls Self-Renewal and Hypoxia Survival in Human Stem/Progenitor Cells of the Mammary Gland Expanded In Vitro as Mammospheres. Stem Cells 25, 807–815.

Saxena, K., and Jolly, M.K. (2019). Acute vs. chronic vs. cyclic hypoxia: Their differential dynamics, molecular mechanisms, and effects on tumor progression. Biomolecules 9, 339.

Saxena, M., Balaji, S.A., Deshpande, N., Ranganathan, S., Pillai, D.M., Hindupur, S.K., and Rangarajan, A. (2018). AMP-activated protein kinase promotes epithelial-mesenchymal transition in cancer cells through Twist1 upregulation. Journal of cell science 131, jcs208314.

Schaffer, B.E., Levin, R.S., Hertz, N.T., Maures, T.J., Schoof, M.L., Hollstein, P.E., Benayoun, B.A., Banko, M.R., Shaw, R.J., and Shokat, K.M. (2015). Identification of AMPK phosphorylation sites reveals a network of proteins involved in cell invasion and facilitates large-scale substrate prediction. Cell metabolism 22, 907–921.

Schroeter, E.H., Kisslinger, J.A., and Kopan, R. (1998). Notch-1 signalling requires ligand-induced proteolytic release of intracellular domain. Nature 393, 382–386.

Scialpi, F., Malatesta, M., Peschiaroli, A., Rossi, M., Melino, G., and Bernassola, F. (2008). Itch self-polyubiquitylation occurs through lysine-63 linkages. Biochemical pharmacology 76, 1515–1521.

Seo, E.J., Kim, D.K., Jang, I.H., Choi, E.J., Shin, S.H., Lee, S.I., Kwon, S.-M., Kim, K.-H., Suh, D.-S., and Kim, J.H. (2016). Hypoxia-NOTCH1-SOX2 signaling is important for maintaining cancer stem cells in ovarian cancer. Oncotarget 7, 55624.

Shin, D.H., Choi, Y.-J., and Park, J.-W. (2014). SIRT1 and AMPK mediate hypoxia-induced resistance of non-small cell lung cancers to cisplatin and doxorubicin. Cancer research 74, 298–308.

Singh, P., Satriano, J., Li, H., and Hallows, K. (2013). HIF-1alpha and AMPK interactions regulate cellular hypoxia adaptation in the subtotal nephrectomy model of CKD (Federation of American Societies for Experimental Biology).

Stylianou, S., Clarke, R.B., and Brennan, K. (2006). Aberrant activation of notch signaling in human breast cancer. Cancer research 66, 1517–1525.

Sundararaman, A., Amirtham, U., and Rangarajan, A. (2016). Calcium-oxidant signaling network regulates AMP-activated protein kinase (AMPK) activation upon matrix deprivation. Journal of Biological Chemistry 291, 14410–14429.

Vacaresse, N., Møller, B., Danielsen, E.M., Okada, M., and Sap, J. (2008). Activation of c-Src and Fyn kinases by protein-tyrosine phosphatase RPTPα is substrate-specific and compatible with lipid raft localization. Journal of Biological Chemistry 283, 35815–35824.

Vaupel, P., Briest, S., and Höckel, M. (2002). Hypoxia in breast cancer: pathogenesis, characterization and biological/therapeutic implications. Wiener Medizinische Wochenschrift 152, 334–342.

Wang, Z., Li, Y., Ahmad, A., Azmi, A.S., Banerjee, S., Kong, D., and Sarkar, F.H. (2010). Targeting Notch signaling pathway to overcome drug resistance for cancer therapy. Biochimica et Biophysica Acta (BBA)-Reviews on Cancer 1806, 258–267.

Wenger, R.H., and Gassmann, M. (1997). Oxygen (es) and the hypoxia-inducible factor-1. Biological chemistry 378, 609–616.

Winder, W., and Hardie, D. (1996). Inactivation of acetyl-CoA carboxylase and activation of AMP-activated protein kinase in muscle during exercise. American Journal of Physiology-Endocrinology And Metabolism 270, E299–E3O4.

Wu, G., Lyapina, S., Das, I., Li, J., Gurney, M., Pauley, A., Chui, I., Deshaies, R.J., and Kitajewski, J. (2001). SEL-10 is an inhibitor of notch signaling that targets notch for ubiquitin-mediated protein degradation. Molecular and cellular biology 21, 7403–7415.

Xi, L., Taher, M., Yin, C., Salloum, F., and Kukreja, R.C. (2004). Cobalt chloride induces delayed cardiac preconditioning in mice through selective activation of HIF-1α and AP-1 and iNOS signaling. American Journal of Physiology-Heart and Circulatory Physiology 287, H2369–H2375.

Xiang, L., Gilkes, D.M., Hu, H., Takano, N., Luo, W., Lu, H., Bullen, J.W., Samanta, D., Liang, H., and Semenza, G.L. (2014). Hypoxia-inducible factor 1 mediates TAZ expression and nuclear localization to induce the breast cancer stem cell phenotype. Oncotarget 5, 12509.

Xing, F., Okuda, H., Watabe, M., Kobayashi, A., Pai, S.K., Liu, W., Pandey, P.R., Fukuda, K., Hirota, S., and Sugai, T. (2011). Hypoxia-induced Jagged2 promotes breast cancer metastasis and self-renewal of cancer stem-like cells. Oncogene 30, 4075–4086.

Yang, C., Zhou, W., Jeon, M.-s., Demydenko, D., Harada, Y., Zhou, H., and Liu, Y.-C. (2006). Negative regulation of the E3 ubiquitin ligase itch via Fyn-mediated tyrosine phosphorylation. Molecular cell 21, 135–141.

Yugawa, T., Handa, K., Narisawa-Saito, M., Ohno, S.-i., Fujita, M., and Kiyono, T. (2007). Regulation of Notch1 gene expression by p53 in epithelial cells. Molecular and cellular biology 27, 3732–3742.

Zadra, G., Batista, J.L., and Loda, M. (2015). Dissecting the dual role of AMPK in cancer: from experimental to human studies. Molecular cancer research 13, 1059–1072.

Zhang, C., Samanta, D., Lu, H., Bullen, J.W., Zhang, H., Chen, I., He, X., and Semenza, G.L. (2016). Hypoxia induces the breast cancer stem cell phenotype by HIF-dependent and ALKBH5-mediated m6A-demethylation of NANOG mRNA. Proceedings of the National Academy of Sciences 113, E2047–E2O56.

Zhang, H., Lu, H., Xiang, L., Bullen, J.W., Zhang, C., Samanta, D., Gilkes, D.M., He, J., and Semenza, G.L. (2015). HIF-1 regulates CD47 expression in breast cancer cells to promote evasion of phagocytosis and maintenance of cancer stem cells. Proceedings of the National Academy of Sciences 112, E6215–E6223.

Zhao, C., Zhang, Q., Yu, T., Sun, S., Wang, W., and Liu, G. (2016). Hypoxia promotes drug resistance in osteosarcoma cells via activating AMP-activated protein kinase (AMPK) signaling. Journal of bone oncology 5, 22–29.

Zheng, X., Linke, S., Dias, J.M., Zheng, X., Gradin, K., Wallis, T.P., Hamilton, B.R., Gustafsson, M., Ruas, J.L., and Wilkins, S. (2008). Interaction with factor inhibiting HIF-1 defines an additional mode of cross-coupling between the Notch and hypoxia signaling pathways. Proceedings of the National Academy of Sciences 105, 3368–3373.

Zhou, G., Myers, R., Li, Y., Chen, Y., Shen, X., Fenyk-Melody, J., Wu, M., Ventre, J., Doebber, T., and Fujii, N. (2001). Role of AMP-activated protein kinase in mechanism of metformin action. The Journal of clinical investigation 108, 1167–1174.

