## Supplementary material for "AMPK-Fyn signaling promotes Notch1 stability to potentiate hypoxia-induced breast cancer stemness and drug resistance": Method and Material

**Materials and Methods:**

**Cell Culture:**

Cell lines MDA-MB-231, BT-474, HCC-1806, MCF-7 (procured from ATCC, VA, USA) were cultured in DMEM (Sigma, MO, USA) with 10% fetal bovine serum (Invitrogen, CA, USA), penicillin and streptomycin. HMLE (kind gift from Dr. Robert Weinberg, MIT, USA) were cultured in DMEM-F12 with growth factors. All cell lines were maintained in standard 5% CO_2_ incubator at 37 °C. AMPKα1/2 DKO MEFs were a kind gift from Dr. Benoit Viollet (INSERM, France). For generation of hypoxic conditions cells were kept in 3% O_2_ in a hypoxia incubator (Eppendorf, Germany).

**Plasmids, transfection and stable cell line generation**

Transfections were carried out using Lipofectamine 2000 (Invitrogen) or Polyethyleneimine (Sigma). EGFP CA CaMKK and pMT2 HA AMPK γ1 R70Q were a kind gift from Dr. Grahame Hardie (University of Dundee, Scotland) and Dr. Lee Witters (Dartmouth College, USA), respectively. pCMV Tag3B AMPK α1 and α2 were a kind gift from Dr. Ronald Evans (Salk institute, USA). pTRIPZ with shRNA against AMPK α2 (2 sequences) were procured from Dharmacon. 12XCSL-Luc was obtained as a kind gift from Dr. Rajan Dighe (IISc, Bangalore). pRK5 Myc-Itch was received as a kind gift from Gerry Melino (University of Leicester, UK), pTriEx-4 Neo DN Fyn K299M and shFYN was a kind gift from Dr. Akash Gulyani (InStem, Bangalore). The shNotch1 (2 sequences) and scrambled constructs were purchased from TransOMIC technology (AL, USA). Stable cells were generated using puromycin (Sigma) selection.

**Pharmacological compounds:**

Pharmacological compounds used were: AMPK activators A-769662 (100 µM) (Abcam, Cambridge, UK), Metformin (500 µM) (Sigma Aldrich), cobalt chloride hexahydrate (150 µM) (Sigma Aldrich), cycloheximide (100µg/mL) (Sigma Aldrich), DAPT (N-[N-(3,5-difluorophenacetyl-L-alanyl)]-S-phenylglycine *t*-butyl ester) (5 µM) (Sigma Aldrich), AMPK inhibitor compound C (10µM) (Calbiochem, Merck Biosciences, CA, USA), Nicotinamide 0.5 mM (Calbiochem, Merck Biosciences), doxycycline (5 µg/mL), MG-132 (5μM), PP2 (10µM) (Calbiochem, Merck Biosciences).

**Statistical analysis**

Statistical analysis was done using GraphPad Prism 5. All data are presented as mean ± SEM. p values < 0.05 were considered as statistically significant. Student’s t-test was used; *** represents p<0.001, **represents p<0.01 and * represents p<0.05. Fischer’s exact test was used for correlation studies.

**Immunoblotting**

Whole cell lysates were prepared with lysis buffer (50 mM Tris pH 7.4, 5 mM EDTA, 250 mM NaCl, 0.5% TritonX, 50 mM NaF, 0.5 mM sodium orthovanadate, 100mM sodium pyrophosphate (Sigma Aldrich), and protease inhibitor (Roche, Switzerland)). 8% SDS PAGE was used to resolve the proteins. α tubulin was used as loading control. Densitometric analysis of the blots was done using Multi Gauge V2.3 (Fujifilm). Intensity of bands were normalised to loading control and represented as fold change over control.

When required, same lysates were run multiple times to assemble individual multi-panel blots; independent loading control (α tubulin) blot was developed for each run.

**Immunoprecipitation**

Cells grown in 90 mm dish were lysed with cold lysis buffer (25mm Tris pH7.4, 1mM EDTA, 150 mM NaCl, 5% glycerol, 0.5 mM sodium orthovanadate, 100mM sodium pyrophosphate, protease inhibitor). 1mg protein from each lysate was estimated and incubated with 1 µg of antibody and 10 µL sepharose G beads (Invitrogen) at 4º C overnight. After incubation the beads were washed with lysis buffer and boiled with Laemmli’s buffer and loaded on SDS-PAGE.

**Primary antibodies**

Antibodies against C-terminal end of Notch1 that recognizes full length (300 kDa) as well as cleaved Notch1 (120 kDa) were purchased from Santa Cruz Biotechnology (Texas, USA), Itch from DSHB, c-Jun antibody and phospho Itch Y420 from Abcam and α tubulin from Calbiochem, Merck. phospho Thr 172 AMPK, phospho Ser 79 ACC, total AMPKα1/2, total AMPKα2, NICD or cleaved Notch1 valine 1744 (gamma-secretase cleavage specific), cyclin D3, Bmi1, Nanog, phospho threonine, phospho serine, acetyl lysine, pan-ubiquitin, K48-linked ubiquitin and phospho Tyr 416 Src family and CA9 were purchased from Cell signalling Technology (MA, USA).

**Immunocytochemistry**

Cells grown in 35 mm dish were fixed with methanol: acetone, permeabilized using 0.2% Triton X 100, blocked with 0.2% fish skin gelatin and incubated with primary antibodies against NICD, Hes5, cy3/Alexa 488 conjugated secondary antibodies (Jackson ImmunoResearch, PA, USA) were used and Hoechst 33342 (1 µg/mL) was used to counterstain and imaged at 20X using Olympus IX71 (Japan). Images were processed using Image Pro Plus (Media Cybernetics).

**Luciferase assay**

MDA-MB-231 cells were seeded in a 60 mm dish were transfected with pGL3-12xCSL luciferase and pRLTK using Lipofectamine 2000 (Invitrogen). The transfected cells were seeded in 12 well plate (1x10^5^ cell per well), and treatment was carried out for 48 h. Cells were harvested for luciferase activity using Dual luciferase assay kit (Promega, WI, USA) as per manufacturer’s instructions and analysed for luciferase activity using TECAN infinite M200 pro plate reader (Switzerland). Firefly luciferase activity was normalised to *Renila* luciferase activity. Data is represented as fold change in relative light units.

**RNA isolation and q-PCR**

Total RNA was isolated from 60 mm dish using TRI reagent (Sigma). cDNA synthesis kit (Applied Biosystems, CA,USA) was used to convert 2 µg RNA to cDNA. q-PCR for Hes1 and Notch1 was performed using SYBR green (Kappa Biosystems) in Eppendorf Mastercycler.

**LCMS/MS**

Liquid Chromatography Mass Spectrometry (LCMS/MS) was carried out to analyse the interactome of Fyn in HEK-293T cells. Protein complexes were isolated from FLAG-tagged expressing HEK-293T stable cells treated with DMSO or A769662 by FLAG immunoprecipitation. 100 µg of the samples was taken for digestion. The sample was diluted with 50 mM NH4HCO3. The sample was treated with 100 mM DTT at 95˚C for 1 h followed by 250 mM IDA at room temperature in dark for 45 min. The sample is then digested with Trypsin and incubated overnight at 37˚C. The resulting sample was vacuum dried and dissolved in 20 µl of 0.1% formic acid in water. After centrifugation at 10000 g, the supernatant was collected into a separate tube.10 µL injection volume was used on BEH C18 UPLC column for separation of peptides. The peptides separated on the column were directed to Waters Synapt G2 Q-TOF instrument for MS and MSMS analysis. The raw data was processed by MassLynx 4.1 WATERS. The individual peptides MSMS spectra were matched to the database sequence for protein identification on PLGS software, WATERS. Fyn interacting partners were identified by processing the raw data through PLGS search engine for protein identification and expression.

**Sphere formation assay**

1x10^5^ cells per 35 mm dish coated with 0.6% agar were seeded in serum-free DMEM-F12 medium with growth factors containing methylcellulose. Sphere forming efficiency was assessed by culturing these cells for 7 days with or without treatment. The primary mammospheres were counted and trypsinised and were allowed to form secondary spheres for another 7 days.

**DNA damage assay**

MDA-MB-231 cells were incubated with 1 µM doxorubicin at 37⁰C for 24 h. After quick PBS wash, cells were fixed using methanol: acetone for 5 min in -20ºC. Cells were incubated with 1:200 anti-pH2A.X antibody at 4⁰C, overnight and subsequently immunofluorescence was carried out. The cells were imaged at 20X and 60X using Olympus IX71. Image processing was done using Image Pro Plus and intensity of puncta was measured.

**MTT assay**

MTT (3-(4,5-Dimethylthiazol-2-yl)-2,5-Diphenyltetrazolium Bromide) is a colorimetric assay that measures cell viability as a function of the redox potential of cells. Actively respiring cells can convert the water soluble MTT to insoluble purple formazan crystals. MTT (1 mg/mL from a 5mg/mL stock) was added to the cells (in 100 µL media) at the end of the experiment and 4 h later the formazan crystals were dissolved in 100 µl of DMSO and the absorbance was measured at 570 nm in a spectrophotometer.

**Immunohistochemistry**

A total of 19 breast cancer tissue samples were obtained from previously untreated grade III invasive ductal carcinoma cases. The paraffin blocks were collected from Kidwai Memorial Institute of Oncology (KMIO; Bangalore, KA, India). Medical Ethics Committee (Institutional Review Board of KMIO) and Institutional Human Ethics Committee (IISc; Bangalore) approved the study. Cleaved Notch1 (Valine 1744) and pACC antibodies were used at 1/100 dilution. IHC was performed as described previously (Sundararaman et al., 2016). Fischer’s exact test was used to test the correlation between NICD and pACC.

### *In-vivo* tumor formation

BT-474 cells (1x10^6^/100 µL) were subcutaneously injected into each flank of the 10 female athymic nude mice. The mice were allowed to form tumors upto 100 mm^3^. Then the mice were randomly divided into 4 groups with 5 mice in each. DMSO was injected as a vehicle control intra-peritoneally and served as the control group. Compound C (2 mg/kg) was intraperitoneally injected into second group for every 4 days. After 24^th^ day the tumors were excised and fixed in 10% formalin for immunohistochemistry for NICD.

**Expression data and analysis:**

TCGA Breast Cancer (BRCA) gene expression RNAseq (IlluminaHiSeq) data and Curated survival data files were downloaded from UCSC Xena (<https://xenabrowser.net/datapages/?cohort=TCGA%20Breast%20Cancer%20(BRCA)&removeHub=https%3A%2F%2Fxena.treehouse.gi.ucsc.edu%3A443>) processed for gene expression in R 3.6.3 (Available on https://www.R-project.org/) (Team, 2013).

**Correlation Analysis**

Pearson’s product moment correlation coefficient, Spearman's rho and their respective significance values between gene pairs across the gene signatures was calculated using cor.test () function in Stats package and correlation matrix generated was visualized using Corrplot (Version 0.84) package in R 3.6.3.(Wei and Simko, 2017)

Codes: (R codes are available on request).

**Hierarchical cluster analysis**

List of signature genes was taken from Molecular Signatures Database GSEA (http://software.broadinstitute.org/gsea/msigdb), AmiGO Gene Ontology Consortium (http://amigo.geneontology.org/amigo)*,* Profiler PCR Array list Qiagen. Expression of genes across breast invasive carcinoma patients in NCBI GEO https://www.ncbi.nlm.nih.gov/geo/query/acc.cgi?acc=GSE40206. Heat maps of combined datasets were generated using GENEPATTERN software.broadinstitute.org/cancer/software/genepattern. We performed Hierarchical cluster analysis and data are categorized into different clusters showing similar expression profiles.

**Mathematical modeling**

Our experimental results indicate that hypoxic conditions enhanced stem-like behavior, through enhanced levels of Notch1, BMI1, and Nanog (Figure 6), OCT4 and SOX2 (data not shown). This stem-like behavior was seen to be sustained in secondary cultures as well, i.e. a hypoxia-persistent signature. Furthermore, literature suggests the presence of mutual activation between the pairs of Notch1-BMI1 (Ohtaka et al., 2017) (Schaller et al., 2010) and OCT4-SOX2 (Chickarmane et al., 2006), and regulation of Notch by Itch (Chastagner et al., 2008). Notch1-mediated upregulation of Sox2 was seen in our observations as well (data not shown), consistent with reported literature (Xiao et al., 2017) (Lee et al., 2016). Integrating this information with our experimental results, we constructed a network and simulated it mathematically using the following set of coupled ordinary differential equations.

$$\frac{dN}{dt}=g_{N}*H^{S+}\left( B,N \right)-k_{N}*N*H^{S-}\left( I,N \right)$$

$$\frac{dB}{dt}=g_{B}*H^{S+}\left( N,B \right)-k_{B}*B$$

$$\frac{dO}{dt}=g_{O}*H^{S+}\left( N,O \right)*H^{S+}\left( S,O \right)-k_{O}*O*H^{S-}\left( I,O \right)$$

$$\frac{dS}{dt}=g_{S}*H^{S+}\left( O,S \right)-k_{S}*S$$

Where N, B, O and S represent Notch1, Bmi1, Oct4 and Sox2. `I’ represents hypoxic signal, i.e., the cumulative effect of hypoxia-AMPK-Fyn-Itch axis (Figure S6D). $g_{N}, g_{B}, g_{O}, g_{S}$ represent the production rates and $k_{N}, k_{B}, k_{O}, k_{S}$ represent the degradation rates of the above nodes respectively. The regulations are modelled using shifted Hill function (Lu et al., 2013):

$$H^{S+|-}\left( X,Y \right)=\frac{X0Y^{n_{X,Y}}}{X0Y^{n}+X^{n}}- \lambda_{X,Y}*\frac{X^{n}}{X0Y^{n}+X^{n}}$$

$\lambda$ represents fold change in Y caused by X. n and X0Y are representatives of the affinity of the interaction between X and Y. When the shifted hill function is included in the production term, these parameters describe the effect of X on the production of Y, whereas in degradation term, they describe the effect of X on the degradation of Y.

The parameters used for simulating the system are listed below:

| Parameter | Value |
| --- | --- |
| $g_{N}$ | 1000* |
| $g_{B}$ | 1000* |
| $g_{O}$ | 1000* |
| $g_{S}$ | 1000* |
| $k_{N}$ | 0.1** |
| $k_{B}$ | 0.1** |
| $k_{O}$ | 0.1** |
| $k_{S}$ | 0.1** |
| $B0N$ | 12600*** |
| $\lambda_{B,N}$ | 3 |
| $n_{B,N}$ | 6 |
| $N0B$ | 18600*** |
| $\lambda_{N,B}$ | 2 |
| $n_{N,B}$ | 6 |
| $I0O$ | 5000*** |
| $\lambda_{I,O}$ | 2 |
| $n_{I,O}$ | 6 |
| $N0O$ | 12400*** |
| $\lambda_{N,O}$ | 4 |
| $n_{N,O}$ | 6 |
| $O0S$ | 18600*** |
| $\lambda_{O,S}$ | 2 |
| $n_{O,S}$ | 6 |
| $S0O$ | 12600*** |
| $\lambda_{S,O}$ | 3 |
| $n_{S,O}$ | 6 |

* - molecules/hr
** - hr^-1^
*** - molecules

This network was then simulated for numerical solutions, and bifurcation diagrams were drawn using MATCONT (Dhooge et al., 2008) . These bifurcation diagrams suggested that as hypoxic response is enhanced, it is possible for cells to switch from a differentiated/non-stem-like state (low Notch1, low OCT4, low SOX2) to a stem-like state (high Notch1, high OCT4, high SOX2) (Figure 6E-G). Interestingly, our model predicts that this switch can be possibly maintained even after the hypoxic/AMPK signal is removed.

Reference:

Chastagner, P., Israel, A., and Brou, C. (2008). AIP4/Itch regulates Notch receptor degradation in the absence of ligand. PloS one *3*.

Chickarmane, V., Troein, C., Nuber, U.A., Sauro, H.M., and Peterson, C. (2006). Transcriptional dynamics of the embryonic stem cell switch. PLoS computational biology *2*.

Dhooge, A., Govaerts, W., Kuznetsov, Y.A., Meijer, H.G.E., and Sautois, B. (2008). New features of the software MatCont for bifurcation analysis of dynamical systems. Mathematical and Computer Modelling of Dynamical Systems *14*, 147-175.

Lee, S.H., Do, S.I., Lee, H.J., Kang, H.J., Koo, B.S., and Lim, Y.C. (2016). Notch1 signaling contributes to stemness in head and neck squamous cell carcinoma. Laboratory investigation *96*, 508.

Lu, M., Jolly, M.K., Levine, H., Onuchic, J.N., and Ben-Jacob, E. (2013). MicroRNA-based regulation of epithelial–hybrid–mesenchymal fate determination. Proceedings of the National Academy of Sciences *110*, 18144-18149.

Ohtaka, M., Itoh, M., and Tohda, S. (2017). BMI1 inhibitors down-regulate NOTCH signaling and suppress proliferation of acute leukemia cells. Anticancer research *37*, 6047-6053.

Schaller, M.A., Logue, H., Mukherjee, S., Lindell, D.M., Coelho, A.L., Lincoln, P., Carson IV, W.F., Ito, T., Cavassani, K.A., and Chensue, S.W. (2010). Delta-Like 4 Differentially Regulates Murine CD4+ T Cell Expansion via BMI1. PloS one *5*.

Sundararaman, A., Amirtham, U., and Rangarajan, A. (2016). Calcium-Oxidant Signaling Network Regulates AMPK Activation Upon Matrix-deprivation. Journal of Biological Chemistry, jbc. M116. 731257.

Team, R.C. (2013). R: A language and environment for statistical computing.

Wei, T., and Simko, V. (2017). R package “corrplot”: Visualization of a Correlation Matrix. Version 084.

Xiao, W., Gao, Z., Duan, Y., Yuan, W., and Ke, Y. (2017). Notch signaling plays a crucial role in cancer stem-like cells maintaining stemness and mediating chemotaxis in renal cell carcinoma. Journal of Experimental & Clinical Cancer Research *36*, 41.
