## Supplementary Figure for "AMPK-Fyn signaling promotes Notch1 stability to potentiate hypoxia-induced breast cancer stemness and drug resistance"

### Slide 1
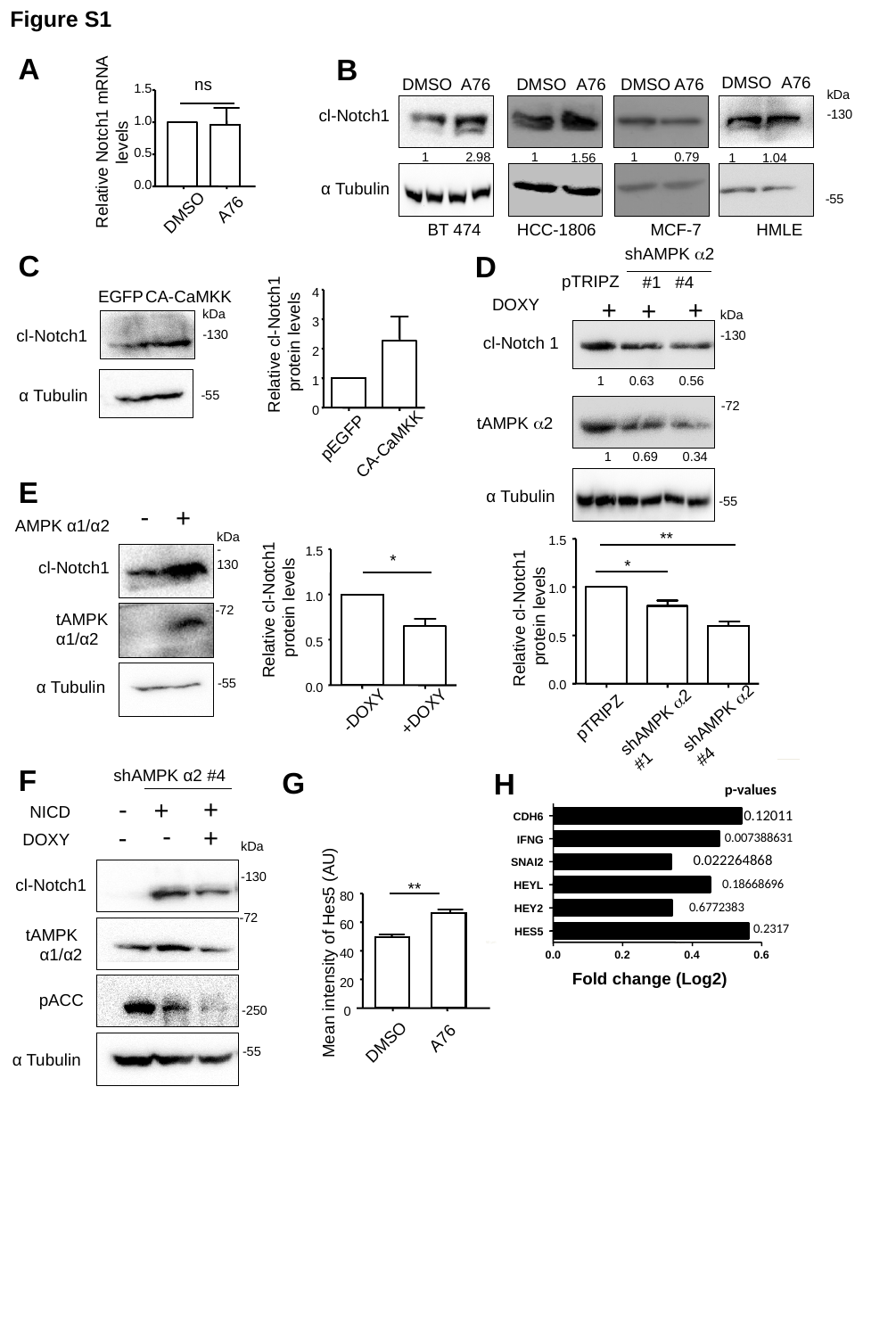

Figure S1
A
B
DMSO
A76
DMSO
A76
DMSO
A76
DMSO
A76
ns
kDa
1.5
cl-Notch1
-130
1.0
Relative Notch1 mRNA levels
1
2.98
1
1
0.79
1.56
1
1.04
0.5
α Tubulin
0.0
-55
A76
DMSO
BT 474
HCC-1806
MCF-7
HMLE
shAMPK α2
C
D
pTRIPZ
#1 #4
 CA-CaMKK
EGFP
4
DOXY
+
+
+
kDa
kDa
3
cl-Notch1
-130
-130
Relative cl-Notch1
protein levels
cl-Notch 1
2
1 0.63 0.56
1
α Tubulin
-55
-72
0
tAMPK α2
pEGFP
CA-CaMKK
1 0.69 0.34
E
α Tubulin
-55
-
+
AMPK α1/α2
**
1.5
*
1.0
Relative cl-Notch1
protein levels
0.5
0.0
shAMPK α2
#4
pTRIPZ
shAMPK α2
#1
kDa
-130
1.5
*
cl-Notch1
Relative cl-Notch1
protein levels
1.0
-72
tAMPK
α1/α2
0.5
-55
α Tubulin
0.0
-DOXY
+DOXY
F
shAMPK α2 #4
G
H
p-values
-
+
+
NICD
0.12011
CDH6
-
-
+
DOXY
0.007388631
kDa
IFNG
0.022264868
**
80
60
 Mean intensity of Hes5 (AU)
40
20
0
A76
DMSO
SNAI2
-130
cl-Notch1
0.18668696
HEYL
0.6772383
HEY2
-72
0.2317
tAMPK
α1/α2
HES5
50 µm
50 µm
0.0
0.2
0.4
0.6
Fold change (Log2)
pACC
-250
-55
α Tubulin

### Slide 2
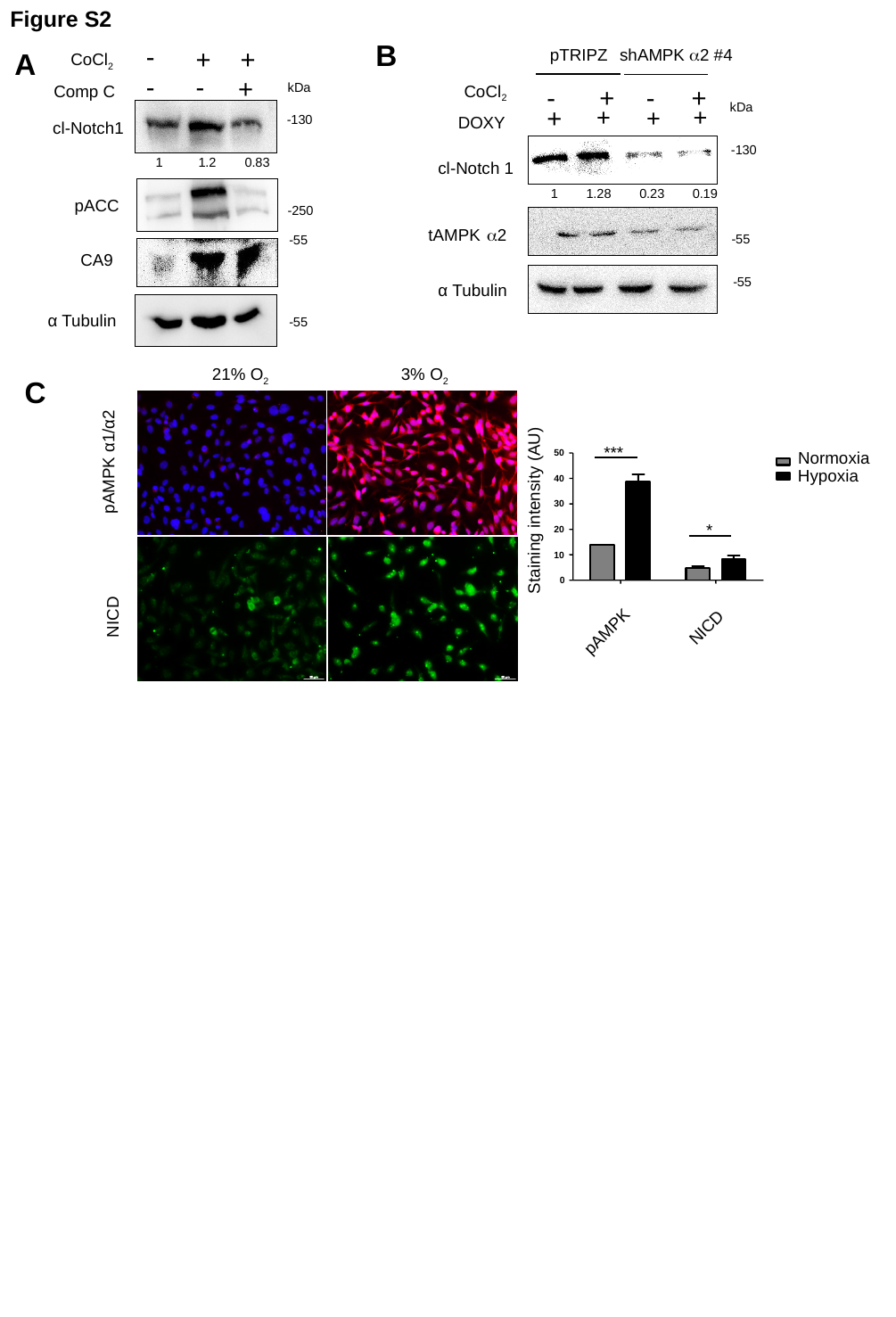

Figure S2
B
-
+
+
shAMPK α2 #4
pTRIPZ
A
CoCl_2_
-
-
+
kDa
CoCl_2_
Comp C
- +
- +
kDa
+
+
+
+
-130
DOXY
cl-Notch1
-130
| 1 | 1.2 | 0.83 |
| --- | --- | --- |
cl-Notch 1
| 1 | 1.28 | 0.23 | 0.19 |
| --- | --- | --- | --- |
pACC
-250
tAMPK α2
-55
-55
CA9
-55
α Tubulin
α Tubulin
-55
3% O_2_
21% O_2_
C
Normoxia
50
Hypoxia
40
30
Staining intensity (AU)
20
10
0
NICD
pAMPK
pAMPK α1/α2
***
*
NICD

### Slide 3
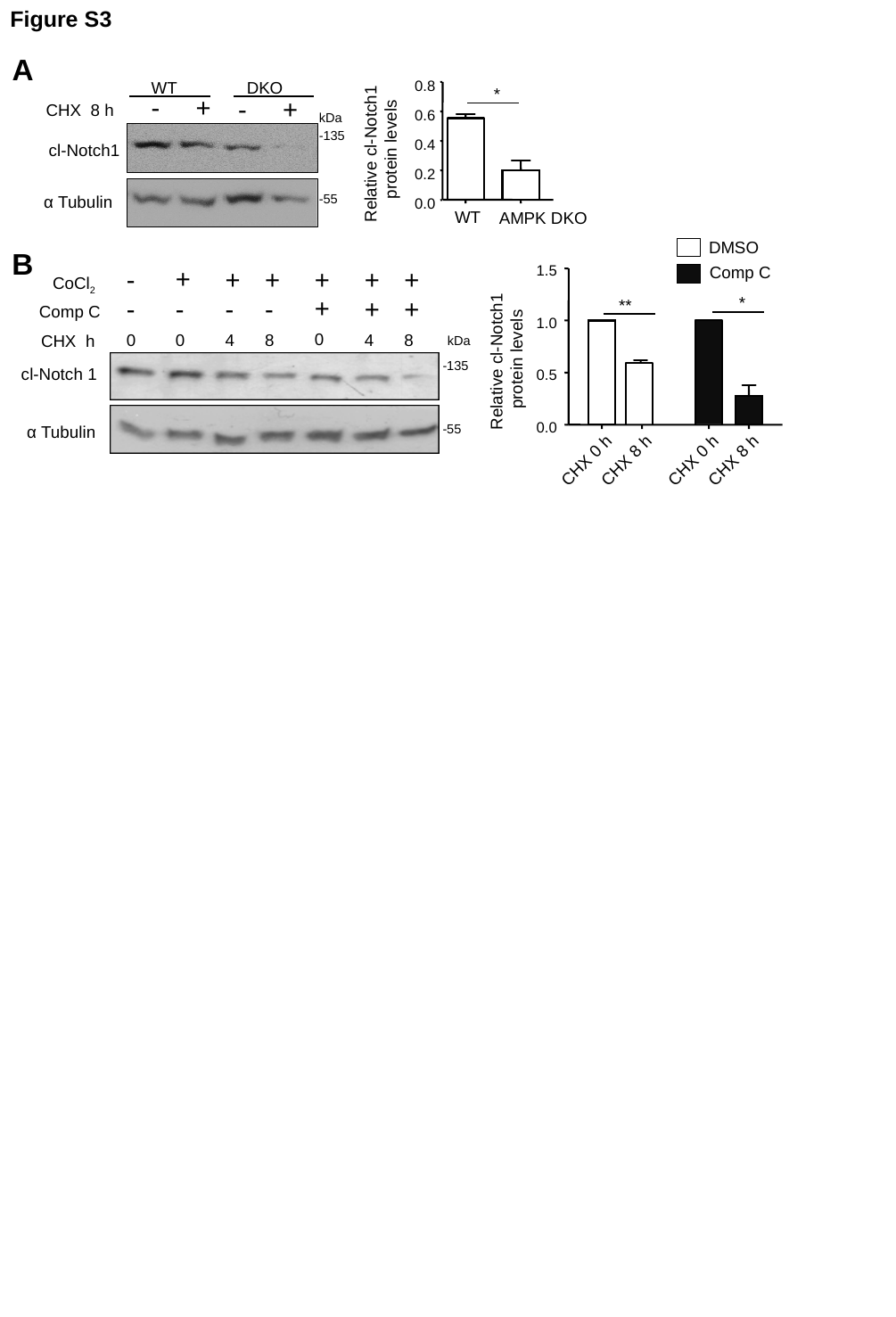

Figure S3
A
WT
 DKO
*
0.8
-
+
-
+
CHX 8 h
kDa
0.6
-135
Relative cl-Notch1
protein levels
cl-Notch1
0.4
0.2
-55
α Tubulin
0.0
 AMPK DKO
WT
DMSO
B
+
-
+
+
+
+
+
CoCl_2_
+
-
-
-
-
+
+
Comp C
0
0
0
4
8
8
4
CHX h
-135
cl-Notch 1
-55
α Tubulin
1.5
Comp C
*
**
1.0
kDa
Relative cl-Notch1
protein levels
0.5
0.0
CHX 0 h
CHX 8 h
CHX 0 h
CHX 8 h

### Slide 4
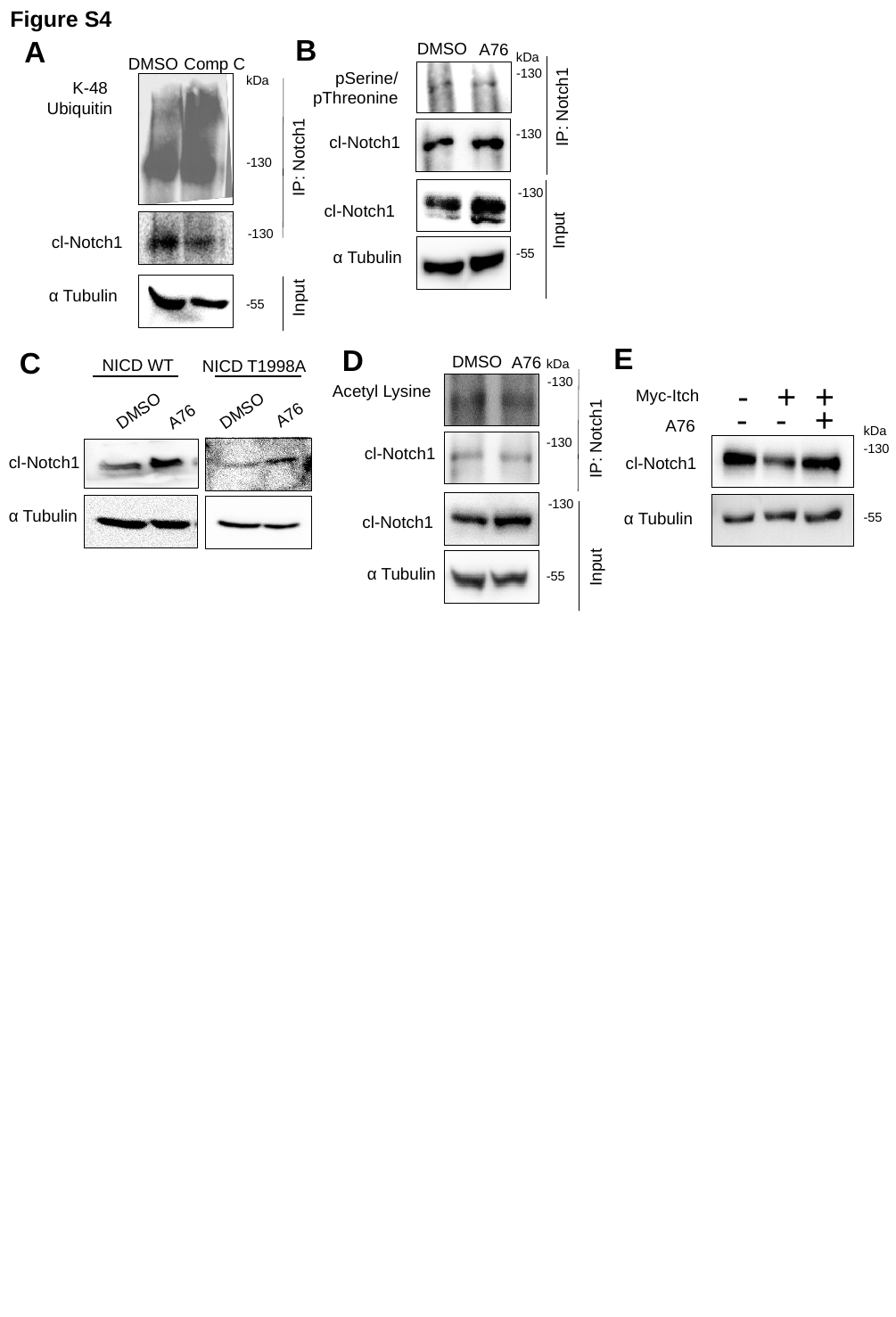

B
DMSO
A76
kDa
-130
pSerine/
pThreonine
-130
cl-Notch1
Input
-130
cl-Notch1
-55
α Tubulin
IP: Notch1
Figure S4
A
DMSO
Comp C
kDa
K-48
Ubiquitin
-130
-130
cl-Notch1
α Tubulin
-55
IP: Notch1
Input
D
DMSO
kDa
-130
Acetyl Lysine
IP: Notch1
-130
cl-Notch1
-130
cl-Notch1
Input
α Tubulin
-55
A76
E
- + +
Myc-Itch
- - +
A76
kDa
-130
cl-Notch1
α Tubulin
-55
C
NICD WT
DMSO
A76
cl-Notch1
α Tubulin
NICD T1998A
DMSO
A76

### Slide 5
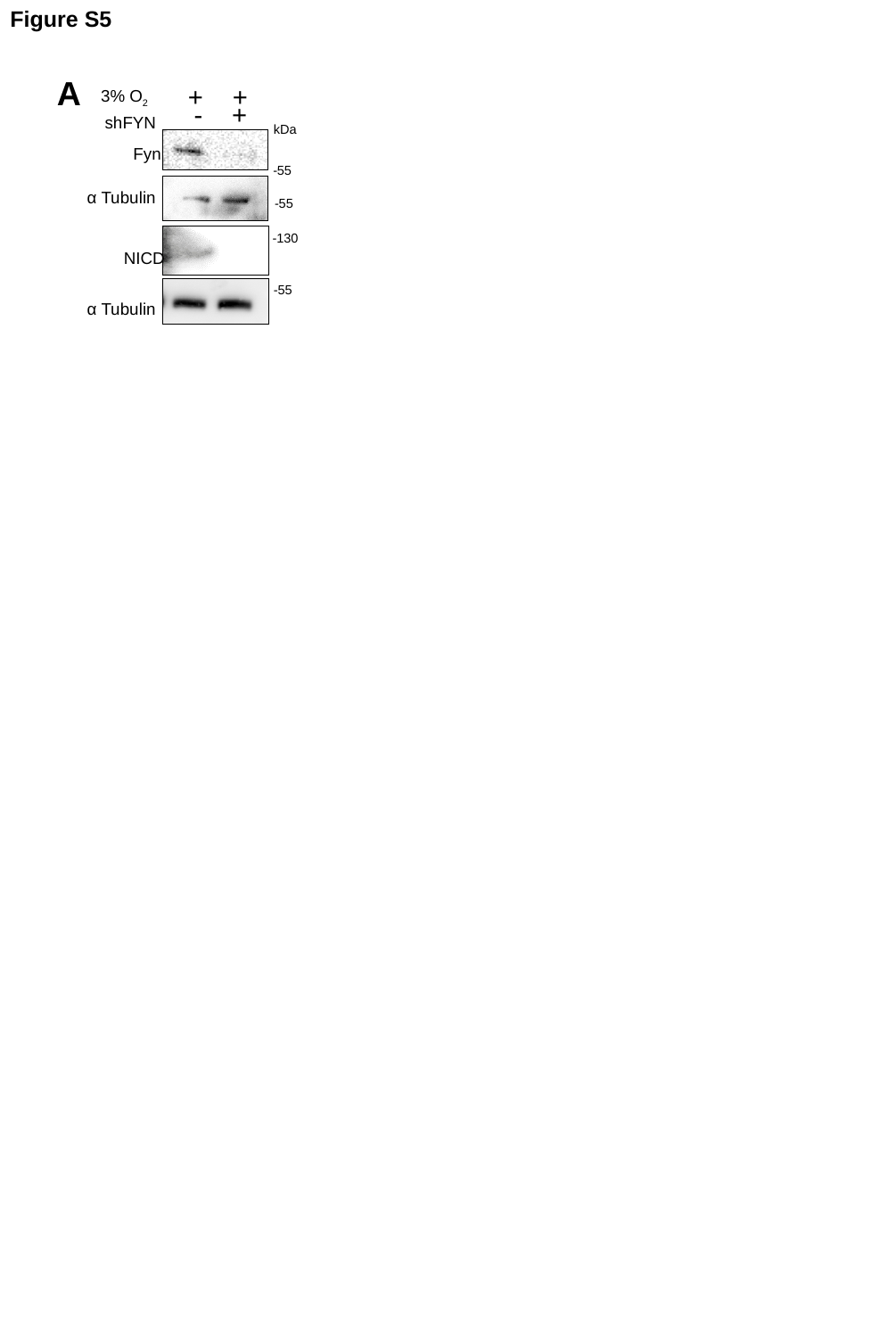

Figure S5
A
+ +
3% O_2_
 - +
shFYN
Fyn
NICD
kDa
-55
-55
-55
-130
α Tubulin
α Tubulin

### Slide 6
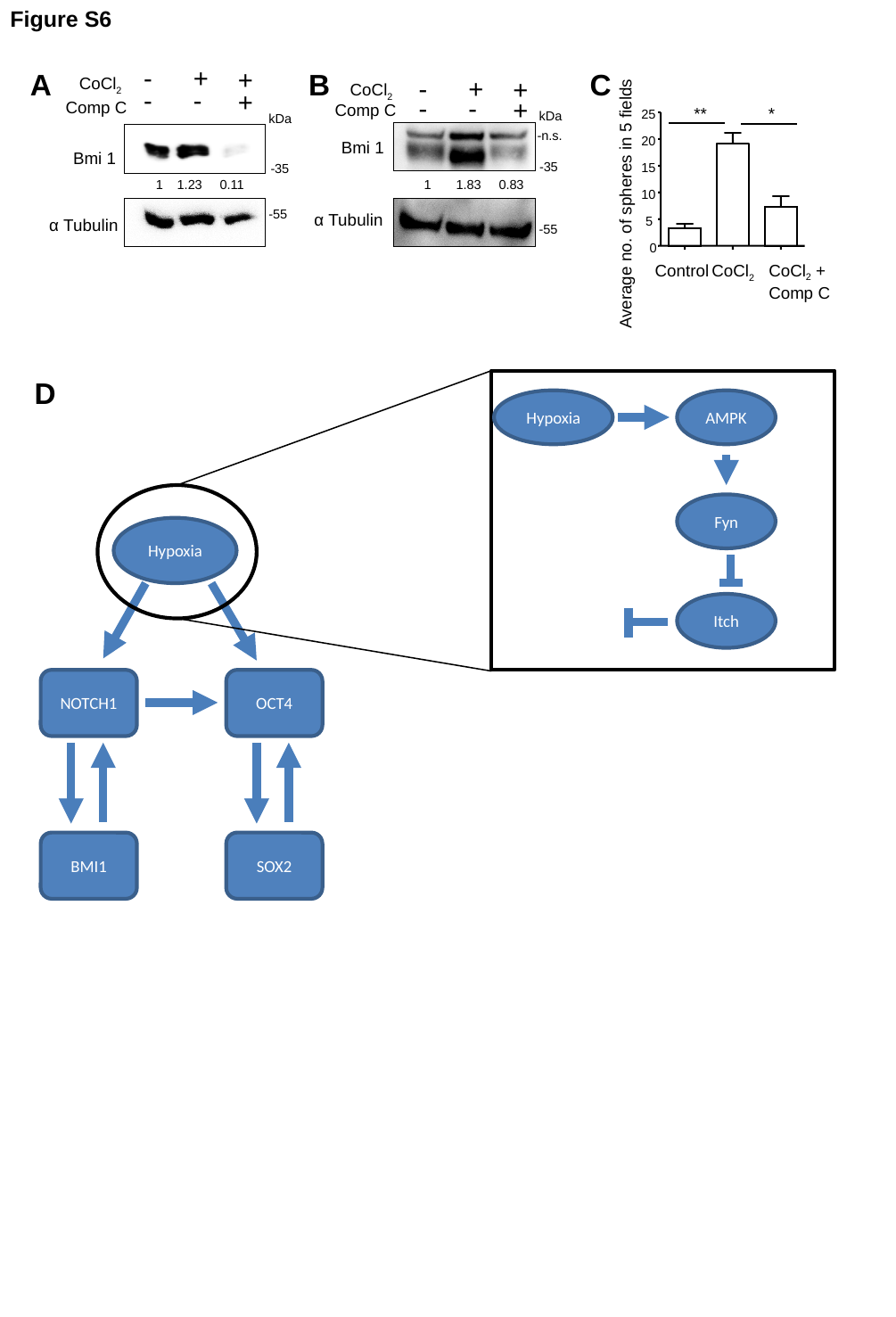

Figure S6
-
+
+
A
B
C
-
+
CoCl_2_
+
CoCl_2_
-
-
**
*
25
20
15
10
Average no. of spheres in 5 fields
5
0
Control
CoCl_2_
CoCl_2_ + Comp C
+
-
-
+
Comp C
 Comp C
kDa
kDa
-n.s.
Bmi 1
Bmi 1
-35
-35
1 1.83 0.83
1 1.23 0.11
-55
α Tubulin
α Tubulin
-55
D
Hypoxia
AMPK
Fyn
Hypoxia
Itch
NOTCH1
OCT4
BMI1
SOX2
-35

### Slide 7
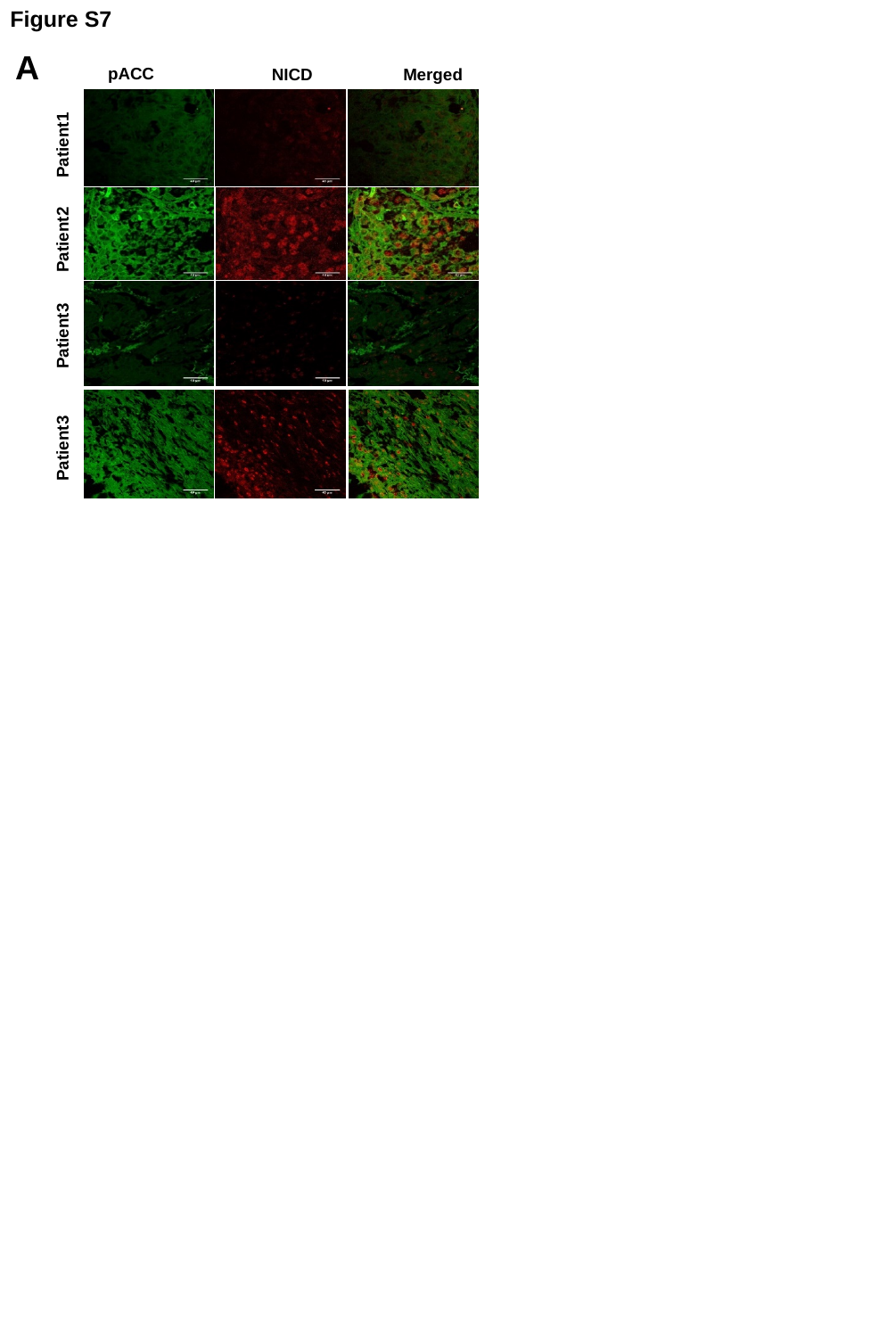

Figure S7
A
pACC
NICD
Merged
Patient1
Patient2
Patient3
Patient3

### Slide 8
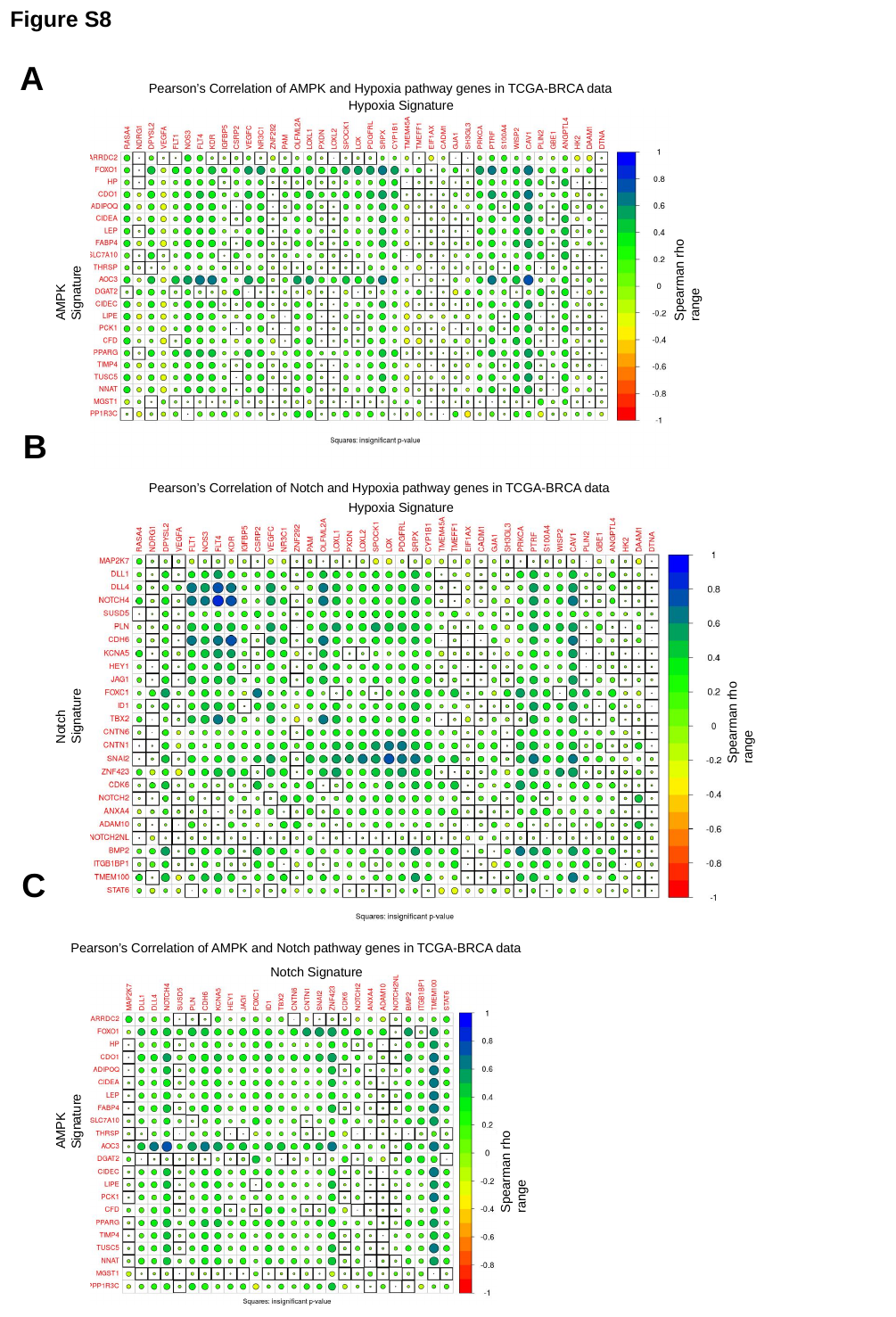

Figure S8
A
B
C
Pearson’s Correlation of AMPK and Notch pathway genes in TCGA-BRCA data
Notch Signature
AMPK Signature
Spearman rho range
Pearson’s Correlation of AMPK and Hypoxia pathway genes in TCGA-BRCA data
Hypoxia Signature
Spearman rho range
AMPK Signature
Pearson’s Correlation of Notch and Hypoxia pathway genes in TCGA-BRCA data
Hypoxia Signature
Spearman rho range
Notch Signature

### Slide 9
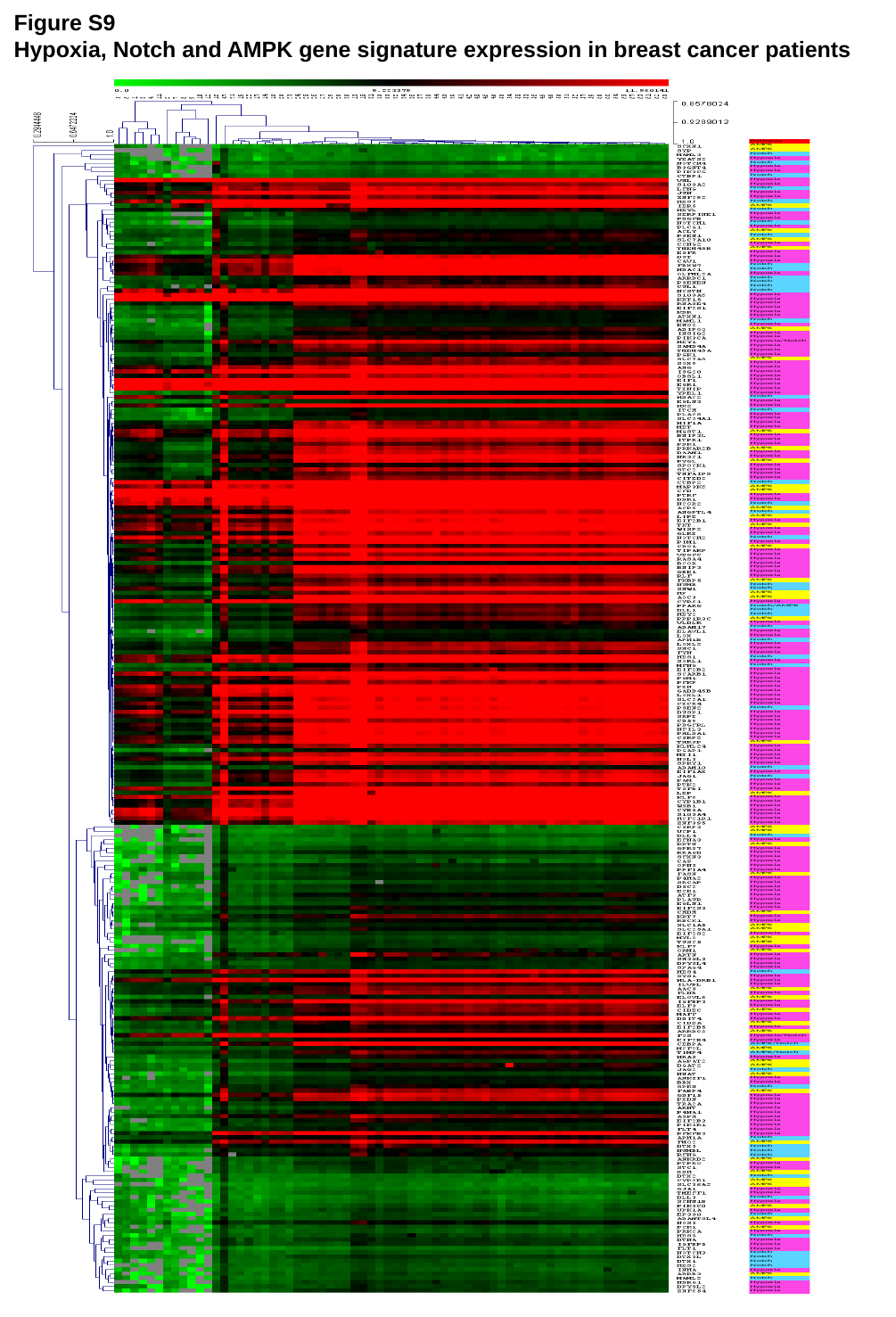

Figure S9
Hypoxia, Notch and AMPK gene signature expression in breast cancer patients
