## Supplementary legends for "AMPK-Fyn signaling promotes Notch1 stability to potentiate hypoxia-induced breast cancer stemness and drug resistance"

**Supplementary Figure legends**

**Figure S1: AMPK regulates cleaved Notch1 levels in breast cancer cells**

(A) MDA-MB-231 cells were cultured in the presence of DMSO or AMPK activator (A76) (100 µM) for 24 h and subjected to RT-PCR analysis for *NOTCH1* and housekeeping gene *GAPDH*; the fold change in transcript levels was determined (n=4). (B) Multiple breast cancer cell lines BT-474, HCC-1806, MCF-7 and immortalized mammary cells HMLE were cultured in the presence of DMSO or A76 (100 µM) for 24 h and were harvested for immunoblotting. The levels of cleaved Notch1 were determined. α Tubulin served as loading control (N=3). (C) MDA-MB-231 cells were transfected with constitutively active form of AMPK-kinase, CaMKKβ, or EGFP as control and were harvested for immunoblotting and probed for cleaved Notch1. Graphs represent the densitometric analysis for quantification of relative amount of indicated protein (n=3). Error bars represent ±SEM. (D) MDA-MB-231 cells stably expressing shRNA against AMPK α2 (#1 and 4) and control pTRIPZ cells were treated with doxycycline (5µg/µl) for 48 h and immunoblotted for AMPK α2 and cleaved Notch1 levels. Graphs represent the densitometric analysis for quantification of relative amount of indicated protein (n=4). Error bars represent ±SEM. (E) AMPK α1/α2 double knockout MEF (DKO) were transfected with AMPK α1 and α2 and were harvested for immunoblotting for cleaved Notch1 and total AMPK α. Graphs represent the densitometric analysis for quantification of relative amount of indicated protein (n=3). Error bars represent ±SEM. (F) HEK-293T cells stably expressing shRNA against AMPK α2 (#4) were transfected with NICD and induced with doxycycline (5µg/µl) for 48 h. Cells were harvested and immunoblotted for cleaved Notch1, AMPKα2 and pACC levels which was compared to uninduced control. Graphs represent the densitometric analysis for quantification of relative amount of indicated protein (n=3). Error bars represent ±SEM. (G) MDA-MB-231 cells were cultured in the presence of DMSO or A76 (100 µM) for 48 h were immunostained for Hes5, 30 cells from each experiment were quantified for Hes5 mean intensity. Graph represent the mean intensity of Hes5 for 3 experiments (n=3). Error bars represent ±SEM. (H) Expression of Notch target genes in microarray analysis of A76-treated MDA-MB-231 cells compared to DMSO treated cells.

**Figure S2:** **AMPK inhibition or knockdown impairs hypoxia-induced cleaved Notch1 generation**

(A) BT-474 cells were cultured in the presence of CoCl_2_ (150 µM) and Comp C (10 µM) for 24 h prior to harvesting. The levels of cleaved Notch1, pACC and CA-9 were determined by immunoblotting (n=3). (B) MDA-MB-231 cells stably expressing shRNA against AMPK α2 (# 4) and control pTRIPZ containing cells were induced with doxycycline (5µg/µl) for 24 h followed by treatment with CoCl_2_ (150 µM) for 24 h and were harvested for immunoblotting and probed for AMPK α2 and cleaved Notch1 levels. (C) MDA-MB-231 cells grown in 21% or 3% O_2_ for 24 h were immunostained for pAMPK and cleaved Notch1 (Valine 1744) levels (Scale bar: 50 μm) (n=3). Graph represent the mean intensity of indicated proteins for 3 experiments. Error bars represent ±SEM.

**Figure S3: AMPK inhibition affects stability of cleaved Notch1 under hypoxia**

(A) Immortalized wild type MEF and AMPK α1/α2 double knockout MEF (DKO) were treated with cycloheximide for 8 h and were harvested for immunoblotting of cleaved Notch1. Graphs represent the densitometric analysis for quantification of relative amount of indicated protein after cycloheximide treatment (n=3). Error bars represent ±SEM. (B) MDA-MB-231 cells were treated with CoCl_2_ along with DMSO or Comp C for 24 h followed by cycloheximide treatment for 4 or 8 h and harvested for immunoblotting for cleaved Notch1 levels. Graphs represent the densitometric analysis for quantification of relative amount of indicated protein (n=3). Error bars represent ±SEM.

**Figure S4: AMPK inhibits Notch1 ubiquitination by modulating its interaction with Itch**

(A) MDA-MB-231 cells were cultured in the presence of DMSO or Comp C (10 µM) for 24 h followed by treatment with MG132 (5 µM) for 2 h prior to harvesting. Cleaved Notch1 was immuno-precipitated followed by immunoblotting and the K-48-linked- ubiquitin levels were determined (n=3). (B) MDA-MB-231 cells were treated with A76 (100 µM) for 24 h and harvested for cleaved Notch1 immuno-precipitation followed by immunoblotting and probed for phospho-serine /threonine levels (n=3). (C) HEK293T cells were transiently transfected with either wild-type NICD or NICD mutant T1998A. These cells were treated with DMSO or A76 (100 µM) for 24 h and harvested for cleaved Notch1 immunoblotting. (D) MDA-MB-231 cells were cultured in the presence of DMSO or A76 (100 µM) for 24 h and harvested for cleaved Notch1 immuno-precipitation followed by immunoblotting and probed for acetylated lysine levels (n=3). (E) BT-474 cells were transfected with Myc tagged Itch plasmid and subsequently treated with DMSO or A76 (100 µM) for 24 h and harvested for immunoblotting of cleaved Notch1 (n=3).

**Figure S5: AMPK mediates Itch phosphorylation in hypoxia**

(A) MDA-MB-231 cells were transiently transfected with control plasmid or shFYN plasmid and immunoblotted for FYN and cleaved Notch1 (Valine 1744) levels.

**Figure S6: Hypoxia-induced AMPK activation promotes stem-like properties in breast cancer cells through Notch signaling**

(A) MDA-MB-231 cells were treated with CoCl_2_ and Comp C (10 µM) as indicated for 48 h prior to harvesting. The levels of stemness proteins Bmi1 were determined by immunoblotting (n=3). (B) BT-474 cells were treated with CoCl_2_ and Comp C (10 µM) as indicated for 48 h prior to harvesting. The levels of stemness proteins Bmi1 were determined by immunoblotting (n=3). (C) MDA-MB-231 cells were treated with CoCl_2_ and Comp C (10 µM) as indicated and then subjected to sphere formation assay. The spheres were counted after 7 days. The graph represents the average number of spheres in 5 fields in two replicates of 3 independent experiments. Error bars represent ±SEM. (D) Regulatory network connecting hypoxic input to stemness through Notch1 has been depicted, integrating our observations with existing literature.

**Figure S7:** **Hypoxia-induced AMPK-Notch1 signaling in vivo**

(A) Representative images from dual fluorescence-immunohistochemical analysis performed on breast cancer patient samples (n=4) to assess cleaved Notch1 (Valine 1744) NICD levels and pACC levels (Scale bar: 40 μm).

**Figure S8:** **Hypoxia-AMPK-Notch gene signature correlations in TCGA Breast cancer dataset:**

**(A-C)** Spearman correlation plots between AMPK-Hypoxia (A), Notch-Hypoxia (B) and AMPK-Notch (C) analyzed using AMPK, Notch and Hypoxia gene signature in TCGA-BRCA (all subtypes) dataset; Spearman correlation value (cor) of each gene pair is represented as the size of the circle and filled with corresponding color from the color palette represented below the ranging from -1(red) to +1(blue). Boxes highlighted by the black squares represent insignificant (p>0.01)

**Figure S9: Hypoxia, Notch and AMPK gene signature expression in breast cancer patients**

Heat map depicting clustering of genes involved in Hypoxia, Notch and AMPK signalling across breast invasive carcinoma patients’ data available in NCBI GEO (GSE40206) data set. Genes of interest are color-coded; AMPK gene signature (yellow), Hypoxia gene signature (pink) and Notch gene signature (blue). Red in the heat map represents highly expressed genes, whereas green represents downregulated genes.
