## Supplementary material for "AMPK-Fyn signaling promotes Notch1 stability to potentiate hypoxia-induced breast cancer stemness and drug resistance": Table1

| Gene_symbols | Signature |
|---|---|
| ARRDC2 | AMPK signature |
| FOXO1 | AMPK signature |
| HP | AMPK signature |
| CDO1 | AMPK signature |
| ADIPOQ | AMPK signature |
| CIDEA | AMPK signature |
| LEP | AMPK signature |
| FABP4 | AMPK signature |
| SLC7A10 | AMPK signature |
| THRSP | AMPK signature |
| AOC3 | AMPK signature |
| DGAT2 | AMPK signature |
| CIDEC | AMPK signature |
| LIPE | AMPK signature |
| PCK1 | AMPK signature |
| CFD | AMPK signature |
| PPARG | AMPK signature |
| TIMP4 | AMPK signature |
| TUSC5 | AMPK signature |
| NNAT | AMPK signature |
| MGST1 | AMPK signature |
| PPP1R3C | AMPK signature |
| RASA4 | Hypoxia |
| NDRG1 | Hypoxia |
| DPYSL2 | Hypoxia |
| VEGFA | Hypoxia |
| FLT1 | Hypoxia |
| NOS3 | Hypoxia |
| FLT4 | Hypoxia |
| KDR | Hypoxia |
| IGFBP5 | Hypoxia |
| CSRP2 | Hypoxia |
| VEGFC | Hypoxia |
| NR3C1 | Hypoxia |
| ZNF292 | Hypoxia |
| PAM | Hypoxia |
| OLFML2A | Hypoxia |
| LOXL1 | Hypoxia |
| PXDN | Hypoxia |
| LOXL2 | Hypoxia |
| SPOCK1 | Hypoxia |
| LOX | Hypoxia |
| PDGFRL | Hypoxia |
| SRPX | Hypoxia |
| CYP1B1 | Hypoxia |
| TMEM45A | Hypoxia |
| TMEFF1 | Hypoxia |
| EIF1AX | Hypoxia |
| CADM1 | Hypoxia |
| GJA1 | Hypoxia |
| SH3GL3 | Hypoxia |
| PRKCA | Hypoxia |
| PTRF | Hypoxia |
| S100A4 | Hypoxia |

| Signatures | gene_list |
|---|---|
| AMPK signatu | 22 |
| Hypoxia | 40 |
| Notch | 26 |

| | |
|---|---|
| WISP2 | Hypoxia |
| CAV1 | Hypoxia |
| PLIN2 | Hypoxia |
| GBE1 | Hypoxia |
| ANGPTL4 | Hypoxia |
| HK2 | Hypoxia |
| DAAM1 | Hypoxia |
| DTNA | Hypoxia |
| MAP2K7 | Notch |
| DLL1 | Notch |
| DLL4 | Notch |
| NOTCH4 | Notch |
| SUSD5 | Notch |
| PLN | Notch |
| CDH6 | Notch |
| KCNA5 | Notch |
| HEY1 | Notch |
| JAG1 | Notch |
| FOXC1 | Notch |
| ID1 | Notch |
| TBX2 | Notch |
| CNTN6 | Notch |
| CNTN1 | Notch |
| SNAI2 | Notch |
| ZNF423 | Notch |
| CDK6 | Notch |
| NOTCH2 | Notch |
| ANXA4 | Notch |
| ADAM10 | Notch |
| NOTCH2NL | Notch |
| BMP2 | Notch |
| ITGB1BP1 | Notch |
| TMEM100 | Notch |
| STAT6 | Notch |
