## Supplementary material for "AMPK-Fyn signaling promotes Notch1 stability to potentiate hypoxia-induced breast cancer stemness and drug resistance": Table2

**Supplementary Table 1: Fyn interacting protein phosphatases identified by MS analysis (**

| Accession | Entry | Descriptior | PLGS Score | Peptides | Coverage (' | peptide.seq |
|---|---|---|---|---|---|---|
| P10586 | PTPRF_HUMAN | Receptor-t | 19.2595 | 23 | 19.9266 | TQRPAMVQTEDQYQLC |
| | | | | | | GSSAGGLQHLVSIR |
| | | | | | | WTEYR |
| | | | | | | DSLLAHSSDPVEMRR |
| | | | | | | QHGQIR |
| | | | | | | FTLTGLKPDTTYDIK |
| | | | | | | EPMDQKR |
| | | | | | | AHTSR |
| | | | | | | TMPVEQVFAK |
| | | | | | | VGGSMLTPR |
| | | | | | | LREMGREK |
| | | | | | | NFRVAAAMK |
| | | | | | | GDGARSKPK |
| | | | | | | IQPLRVQR |
| | | | | | | LPVPSKQHGQIR |
| | | | | | | LVNIMPYELTRVCLQPIR |
| | | | | | | VLAFTAVGDGPPSPTIQ' |
| | | | | | | SANYTCVAISSLGMIEAT |
| | | | | | | VAAAMKTSVLLSWEVPI |
| | | | | | | ACNPLDAGPMVVHCSA |
| | | | | | | IVTTTGAVPGRPTMMIS |
| | | | | | | YASSPYSDEIVVQVTPAC |
| | | | | | | SGALQIESSEESDQGKYE |
| P28827 | PTPRM_HUMAN | Receptor-t | 26.1367 | 11 | 10.0551 | DAPLKEIK |
| | | | | | | LWLQGIDVRDAPLK |
| | | | | | | TVVHCLNGGGR |
| | | | | | | TFAVEKR |
| | | | | | | DTPVSK |
| | | | | | | SNSPPGLLNVYVK |
| | | | | | | ETMSSTR |
| | | | | | | TLNMVTPTLRVEDCSIAI |
| | | | | | | VNMVQTEEQYVFIHDAI |
| | | | | | | SRQITIR |
| | | | | | | MVWHENTASIIMVTNL' |
| Q9H3S7 | PTN23_HUMAN | Tyrosine-pı | 11.1011 | 12 | 7.5795 | VAALLER |
| | | | | | | REWAK |
| | | | | | | LLREMMAK |
| | | | | | | VDAAEGR |
| | | | | | | AELAEVR |
| | | | | | | KKPPPRPTAPKPLLPR |
| | | | | | | GRSIAIAR |
| | | | | | | LELLRQNAVR |
| | | | | | | SIAIARCYSLK |
| | | | | | | MEAVPRMPMIWLDLK |

| | | | | | | |
|---|---|---|---|---|---|---|
| | | | | | | STETHVERVLSLQFR |
| | | | | | | ELIQKDDITASLVTTDHS |
| Q15262 | PTPRK_HUMAN | Receptor-t | 11.0137 | 11 | 10.7019 | IAEIQAR |
| | | | | | | TDQDLYR |
| | | | | | | GYNEIR |
| | | | | | | TFTLER |
| | | | | | | IAEIQARR |
| | | | | | | SFLKLILQVEK |
| | | | | | | GLNPGTLNILVRVNK |
| | | | | | | FMDMLPPDRCLPFLITID |
| | | | | | | TQCVRIATK |
| | | | | | | MTSGSWTETHAVNAPT |
| | | | | | | SYADQSTLHAEDPLSITF |
| O14522 | PTPRT_HUMAN | Receptor-t | 9.734 | 8 | 10.1319 | ASLAALALSLLLR |
| | | | | | | YGNIISYDHSR |
| | | | | | | TVVHCLNGGGR |
| | | | | | | MIWQENSASIVMVTNL |
| | | | | | | VADLLQHITQMKR |
| | | | | | | MIWQENSASIVMVTNL |
| | | | | | | VRPEDCSIGLLPRNHDK |
| | | | | | | AQSAAGGCSFDEHYSN |

**(A769662 treatment)**

:YR

VKTQQGVPAQPADFQAEVESDTR
'AQVTVK
DSYK
.GVGR
TTAMNTALLQWHPPKELPGELLGYR
QQQEEPEMLWVTGPVLAVILIILIVIAILLFKR
:CVATNSAGTR

LLPR
ILEACLCGDTSVPASQVR

VEVGRVK

EMK

GESSNYINAALMDSYR

YKLWHLDPDTEYEIR
MDQHNFSPR

VEVGR

VEVGRVK

GYSVALGTNGFTWEQINTWEKPMLDQAVPTGSFMMVNSSGR
