## Supplementary material for "AMPK-Fyn signaling promotes Notch1 stability to potentiate hypoxia-induced breast cancer stemness and drug resistance": Table3

Table 2: Fyn interacting protein phosphatases identified by MS analysis (DMSO treatment)

| Accession | Entry | Descriptior | PLGS Score | Peptides | Coverage (' | peptide.seq |
|---|---|---|---|---|---|---|
| P54829 | PTN5_HUN | Tyrosine-pı | 46.4739 | 6 | 14.6903 | APPLLHLVR |
| | | | | | | QLSHQSPE |
| | | | | | | TTCQLR |
| | | | | | | ENHAADDSEGGALDMCCSER |
| | | | | | | ALDPFLLQAEFFEIPMNFVDPK |
| | | | | | | QSVSRQPSFTYSEWMEEK |
| P08575 | PTPRC_HU | Receptor-t | 20.0599 | 14 | 13.727 | LFTAK |
| | | | | | | ATVIVMVTR |
| | | | | | | INQHK |
| | | | | | | EQAEGSEPTSGTEGPEHSVNGPASP/ |
| | | | | | | FTNASK |
| | | | | | | HELEMSK |
| | | | | | | DLQYSTDYTFK |
| | | | | | | ELISMIQVVKQK |
| | | | | | | YVLSLHAYIIAKVQR |
| | | | | | | ATVIVMVTR |
| | | | | | | VNAFSNFFSGPIVVHCSAGVGR |
| | | | | | | LPQKNSSEGNK |
| | | | | | | YINASFIMSYWKPEVMIAAQGPLK |
| | | | | | | HELEMSKESEHDSDESSDDDSDSEEP |
| Q92932 | PTPR2_HU | Receptor-t | 56.5868 | 15 | 19.1133 | EQFEFALTAVAEEVNAILK |
| | | | | | | QPAEVR |
| | | | | | | HSSQHR |
| | | | | | | GVPSSSR |
| | | | | | | NLQTNETR |
| | | | | | | VSANVQNVTTEDVEK |
| | | | | | | ATVDNKDK |
| | | | | | | ISSVSSQFSDGPIPSPSAR |
| | | | | | | EIDIAATLEHLRDQRPGMVQTKEQFE |
| | | | | | | LRVALQK |
| | | | | | | VSANVQNVTTEDVEKATVDNK |
| | | | | | | EDLLPRTLGQLQPDELSPK |
| | | | | | | NRSLAVLTYDHSR |
| | | | | | | VLLKAENSHSHSDYINASPIMDHDPR |
| | | | | | | WPSPLGDSEDPSSTGDGAR |
| P23467 | PTPRB_HU | Receptor-t | 21.4839 | 26 | 16.4747 | IQILTVSGGLFSK |
| | | | | | | VTSYEVQLFDENNQK |
| | | | | | | DLLLIHK |
| | | | | | | SFNIK |
| | | | | | | TVPLAVLQLR |
| | | | | | | DVLISK |
| | | | | | | ILQQLDSK |
| | | | | | | CAENPNSNSK |

| | | | | | |
|---|---|---|---|---|---|
| | | | | | NSSSVK |
| | | | | | TVVLQTDPLPPAR |
| | | | | | ELVPGR |
| | | | | | VTNDGSLTSLKVK |
| | | | | | DLTLRNR |
| | | | | | VANLEANNNGRMR |
| | | | | | TAPMEVSNLKVTNDGSLTSLK |
| | | | | | TVPASVSHLR |
| | | | | | TVPSSVSGVTVNNSGR |
| | | | | | SFSVYTNGSTVPSPVKDIGISTK |
| | | | | | TAPSPPSLMSFADIANTSLAITWK |
| | | | | | TVPASVSHLR |
| | | | | | SGSLYSVVVTTVSGGISSRQVVVEGR |
| | | | | | MRSLVVSWSPPAGDWEQYR |
| | | | | | LSNVDDDPCSDYINASYIPGNNFRR |
| | | | | | LYSVTVTTKSGQYEANEQGNGR |
| | | | | | NTSEPATTKQHK |
| | | | | | LYTVTITTRSGK |
| Q92729 | PTPRU_HU | Receptor-t | 9.5372 | 8 | 7.6072 QSGALVPAAGVR |
| | | | | | FLATFPLAAVSR |
| | | | | | GLAFAEIQAR |
| | | | | | KEIEYR |
| | | | | | RPLEVSQR |
| | | | | | AACKESK |
| | | | | | SNHFIATQGPKPEMVYDFWRMVW |
| | | | | | SAAFIVTLHPLQSTTPDFWR |

ALNQGS

PSK

EFALTAVAEEVNAILK

[illegible]

QEHCSSIVMITK
